# MAPK/MAK/MRK overlapping kinase (MOK) controls microglial inflammatory/type-I IFN responses via Brd4 and is involved in ALS

**DOI:** 10.1101/2023.01.23.524851

**Authors:** Jesús A. Pérez-Cabello, Lucía Silvera-Carrasco, Jaime M. Franco, Vivian Capilla-González, Alexandros Armaos, María Gómez-Lima, Raquel García-García, Xin Wen Yap, M. Magdalena Leal-Lasarte, Deepti Lall, Robert H. Baloh, Salvador Martínez, Yoshihiko Miyata, Gian G. Tartaglia, Ritwick Sawarkar, Mario García-Dominguez, David Pozo, Cintia Roodveldt

## Abstract

Amyotrophic lateral sclerosis (ALS) is a fatal and incurable neurodegenerative disease affecting motor neurons and characterized by microglia-mediated neurotoxic inflammation whose underlying mechanisms remain incompletely understood. In this work we reveal that MAPK/MAK/MRK overlapping kinase (MOK), with unknown physiological substrate, displays an immune function by controlling inflammatory and type-I IFN responses in microglia which are detrimental to primary motor neurons. Moreover, we uncover the epigenetic reader bromodomain-containing protein 4 (Brd4) as the first molecule regulated by MOK, by promoting Ser^492^-phospho-Brd4 levels. We further demonstrate that MOK regulates Brd4 functions by supporting its binding to cytokine gene promoters, therefore enabling innate immune responses. Remarkably, we show that MOK levels are increased in ALS spinal cord, particularly in microglial cells, and that administration of a chemical MOK-inhibitor to ALS model mice is able to modulate Ser^492^-phospho-Brd4 levels, suppress microglial activation and modify disease course, indicating a pathophysiological role of MOK kinase in ALS and neuroinflammation.

## Introduction

Amyotrophic lateral sclerosis (ALS) is a fatal adult-onset neurodegenerative disease in which motor neurons in the cortex, brain stem and spinal cord gradually degenerate and die off (1, 2). A typical characteristic of ALS and most other neurodegenerative diseases is neuroinflammation, which is associated with disease progression and spreading (1). In particular, several studies have shown that microglia, the main immunocompetent cells in the central nervous system (CNS), become activated and neurotoxic and thereby contribute to motor neuron loss and disease progression (3). Still, the mechanisms driving microglia activation and neurotoxicity in ALS remain incompletely understood, and key molecular mediators along the activated signalling pathways that may serve as new therapeutic targets or disease biomarkers need to be identified (4–6). A powerful emerging view poses that dysregulated neuroinflammatory responses in these disorders may be caused by disruption at the signalling-epigenetics-transcriptional axis level (7, 8).

A hallmark of ALS and other related disorders is the intracellular accumulation of protein inclusions containing misfolded TDP-43 (transactive response DNA-binding protein-43) and in certain cases, RNA-binding protein FUS or SOD1 (superoxide dismutase 1) proteins. TDP-43 -a multifunctional protein with RNA/DNA binding activities and physiological localization in the cell nucleus-has been found to display functional abnormalities, including its identification as a major component of cytoplasmic intraneuronal and glial inclusions, in virtually all ALS patients (9, 10) and various ALS animal models (11, 12). Thus, TDP-43 is currently thought to represent a common molecular hub in these disorders, where different pathogenic mechanisms may converge and finally result in neurotoxic cellular outcomes (13).

Recently, we reported that MAPK/MAK/MRK overlapping kinase (MOK), a Ser/Thr kinase that belongs to the MAPK superfamily (14), interacts and co-localizes with cytoplasmic TDP-43 inclusions in microglia resulting from exposure of cells to exogenous TDP-43 aggregates (15). MOK is an atypical Ser/Thr signalling kinase, found both in the cytosol and nucleus of different cell types and with a demonstrated function in cilium growth regulation in renal epithelium (16). Up to now, no physiological substrate or downstream-regulated target has been identified for this kinase, and no function or molecular mechanism in relation to immunity or the CNS has been identified.

In this study, we uncover a role of MOK kinase in controlling inflammatory and type-I IFN responses in microglia via a Brd4-dependent mechanism and report that MOK-mediated immune functions are dysregulated, and actively participate, in ALS pathophysiology. This represents both a novel signalling pathway in neuroinflammation and a newly discovered underlying mechanism of ALS-associated immune dysregulation.

## Results

### MOK-mediated mechanisms are mobilized in microglial cells under TDP-43 aggregation

To evaluate whether MOK is engaged in cellular processes activated in the context of TDP-43 aggregation, we started by carrying out SLAM-Seq analysis to report on immediate changes in RNA synthesis (17) from SIM-A9 microglial cells exposed to TDP-43-aggregates (TDP43) or sham controls (Sham), with or without pre-treatment with AG2P145D/Comp13 (C13) MOK inhibitor (18) (**Supplem. Figure 1a, b**). C13 is an unusually specific inhibitor based on the structural, virtual uniqueness of a cysteine gatekeeper in MOK’s active site (18). The results showed a clear difference in the transcriptional changes caused by MOK-inhibition for a number of genes comparing the various conditions. Remarkably, pre-treatment with C13 revealed a dramatically different impact in the number of the differentially expressed genes (DEGs) (**Figure 1a, Supplem. Figure 1a**) and their predicted interaction networks (19) under TDP-43 aggregation compared to control (sham)-treated cells (**Figure 1a**). Interestingly, 45% of the DEGs resulting from exposure to TDP-43 aggregates were counter-regulated by C13 pre-treatment (**Figure 1b, Supplem. Figure 1b**), suggesting a possible participation of MOK in microglial responses elicited by TDP-43 aggregates.

**Figure 1.**
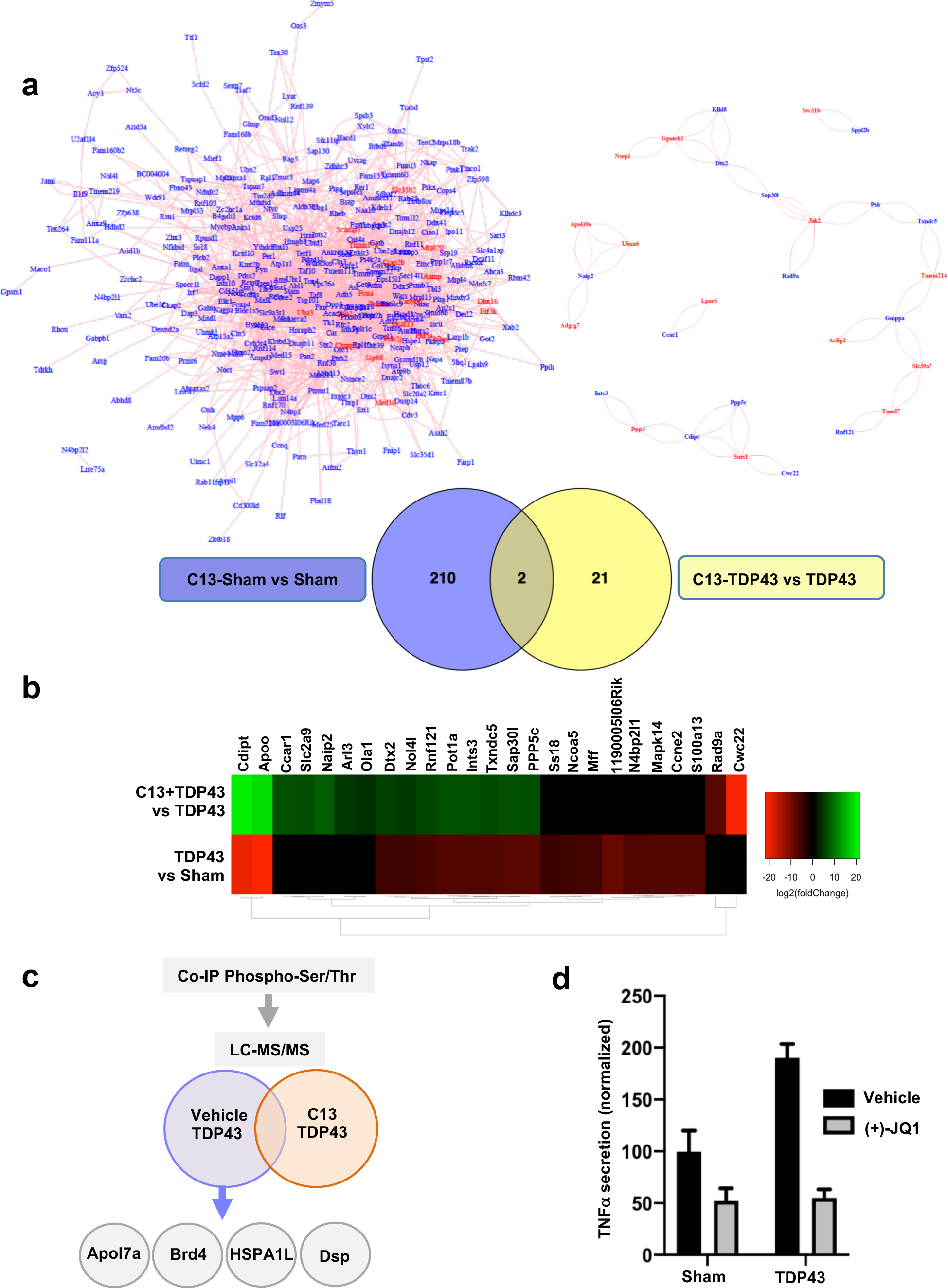
MOK-mediated mechanisms occur in microglial cells upon exposure to TDP-43 aggregates. **a, bottom)** Venn diagrams showing the number of DEGs from analyzed SLAM-Seq data obtained with SIM-A9 cells exposed to 5 μg/mL TDP-43 aggregates (TDP43) or sham aggregates (Sham) for 4 h, pre-treated for 1 h with 10 μM C13 (or DMSO as vehicle). Results are from three independent experiments (N=3). Indicated are the number of DEGs for ‘C13-Sham vs. Sham’ and ‘C13-TDP43 vs. TDP43’ comparisons (pAdj.<0.05 and pAdj.<0.1, respectively). **a, top**) Predicted network analysis based on the two sets of identified DEGs by using GeneMANIA^19^. Blue labels correspond to top DEGs and red labels, to inferred genes. **b**) Heatmap representing the relative expression profiles (DEGs with pAdj.<0.05) comparing ‘TDP43 vs. Sham’ and ‘C13-TDP43 vs. TDP43’ treatments. The differential regulation in gene expression changes between both comparisons indicates an effect of MOK inhibitor C13 in transcriptional profile of microglial cells upon exposure to TDP-43 aggregates. **c**) Schematic representation of LC-MS/MS analysis of eluates from anti-phospho-Ser/Thr immunoprecipitation assays of lysates from primary microglial cells exposed to 5 μg/mL TDP-43 aggregates, pre-treated for 1 h with 10 μM C13 (or DMSO as vehicle). **d**) Determination of TNFα by ELISA from primary microglial cells exposed to 5 μg/mL TDP-43 aggregates (TDP43) or sham aggregates (Sham) overnight, pre-treated for 1 h with 10 μM (+)-JQ1 or DMSO (vehicle). Shown values were normalized to Sham control. Data are mean −/+ S.D. (N=2).

Next, to identify proteins whose phosphorylation state could be regulated by MOK kinase in the ALS cellular model, we used immunoprecipitation against phospho-Ser/Thr-containing proteins from primary microglial cells exposed to TDP-43 aggregates, pre-treated or not with C13. By subsequent LC-MS/MS analyses of both eluates, we identified several proteins in both samples, a few of them exclusively in the absence of MOK inhibitor pre-treatment, including desmoplakin (Dsp), heat-shock protein 1-like protein (HSPA1L), bromodomain-containing protein 4 (Brd4) and apolipoprotein L 7a (Apol7a) (**Figure 1c**). Interestingly, Brd4 -a member of the BET bromodomain family-has been recently involved in innate immune functions, including microglial responses (20–22). To verify the participation of Brd4 in TDP-43-induced microglial activation, we assessed the impact of the Brd4 inhibitor (+)-JQ1 in TNFα secretion by ELISA (**Figure 1d**). Remarkably, the aggregates-elicited cytokine levels were largely suppressed by pre-treatment with (+)-JQ1. Taken together, these results indicate that MOK is functionally mobilized in microglial cells under TDP-43 aggregation and may regulate downstream phosphorylation of proteins, including Brd4, involved in microglial activation and immune responses.

### MOK controls microglial responses by mediating inflammatory and type-I IFN pathways

Given that exposure of microglial cells to pathology-associated TDP-43 species had been found to activate the NLRP3 inflammasome by our group and by others (15, 23, 24), we sought to assess the impact of MOK inhibition in inflammasome activation by using microglial cells exposed to TDP-43 or sham aggregates. Remarkably, the higher levels of IL-1β and IL-18 inflammasome-dependent cytokines induced by TDP-43 aggregates were significantly suppressed by pre-treatment with the MOK-inhibitor C13 (**Supplem. Figure 1c,d**).

Next, in order to investigate a potential role of MOK in mediating the general inflammatory response in microglia, we assessed TNFα, IL-6 and IL-1β cytokine secretion levels in culture supernatants from lipopolysaccharide (LPS)-stimulated primary microglial cells (5 h or 16 h post-stimulation), either pre-treated or not with C13 (**Figure 2a**). Remarkably, C13-pre-treatment significantly reduced secretion of all three proinflammatory cytokines otherwise induced by LPS alone. This result was also observed for SIM-A9 microglial cells, in which pre-treatment with C13 led to lower TNFα secreted levels after LPS stimulation (**Supplem. Figure 2a**). Furthermore, Western blot analyses evidenced lower Ser^536^ phosphorylation of p65 signal upon LPS stimulation for C13-pre-treated cells, pointing to a role of MOK in NF-κB pathway in the inflammatory response (**Supplem. Figure 2b**). Next, to elucidate the effect of C13-treatment on LPS-stimulated primary microglia at a transcriptional level, RNA-Seq was carried out with RNA isolated from primary microglial cells after 5h-stimulation with LPS, subjected or not to pre-treatment with C13. Based on the 158 differentially expressed genes (DEGs) identified, we used Ingenuity Pathway Analysis (IPA) software enabling to predict canonical pathways, upstream regulators, functions and networks, among others (**Figure 2b**). Notably, IPA revealed a number of significantly inhibited (IRF3/7, PML and STAT1) and activated (CEBPB, NFE2L2/Nrf2 and STAT3) upstream regulators as a result of C13 pre-treatment.

**Figure 2.**
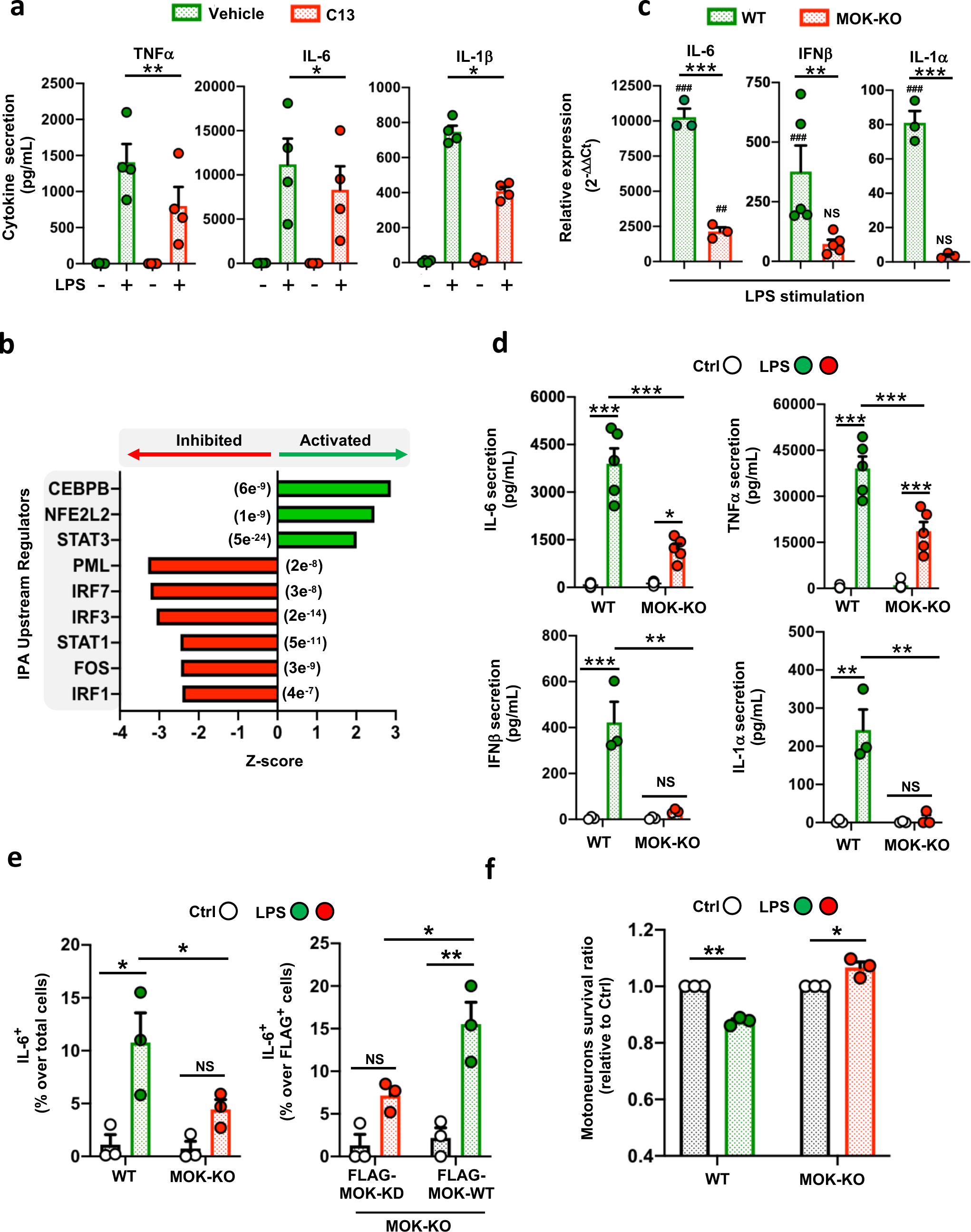
MOK positively regulates microglial responses, including inflammatory and type-I IFN pathways. **a)** Quantification of pro-inflammatory cytokines by ELISA in supernatants from primary microglial cells stimulated with 1 μg/mL LPS for 5 h (TNFα) or 24 h (IL-6, IL-1β) after pre-treatment with 10 μM C13 (or DMSO) for 1 h (N=4). One-way ANOVA followed by Tukey post-hoc test. **b**) Top predicted upstream regulators by Ingenuity Pathway Analysis (IPA) from 158 DEGs (C13-treated vs. untreated, pAdj<0.05) identified by RNA-Seq studies from primary microglial cells stimulated with 1 μg/mL LPS for 5 h. The P-values of overlap are shown in parentheses. **c**) Assessment of gene expression levels for a set of key cytokines by qRT-PCR from WT and MOK-KO SIM-A9 cells stimulated with 1 μg/mL LPS for 5 h (N=3). Data represent fold-changes relative to unstimulated WT cells (values were 1 or lower for unstimulated WT and MOK-KO controls; not shown). ^#^P<0.05, ^##^P<0.01 and ^###^P<0.001 vs. unstimulated controls. One-way ANOVA followed by Tukey post-hoc test. **d**) Quantification of TNFα, IL-6, IFNβ and IL-1α levels by ELISA in supernatants from WT and MOK-KO SIM-A9 cells stimulated with 1 μg/mL LPS for 5 h or overnight for IL-1α (N=5 for IL-6, TNFα and N=4 for IFNβ, IL-1α). One-way ANOVA followed by Tukey post-hoc test. **e**) Flow cytometry quantification of IL-6^+^ cells in WT and MOK-KO SIM-A9 cells, previously stimulated with 1 μg/mL LPS for 5 h (left). Rescue of phenotype in MOK-KO cells resulting from cell transfection with FLAG-MOK-WT or FLAG-MOK-KD constructs and stimulation with 1 μg/mL LPS for 5 h (right) (N=3). Student’s t-test performed between shown groups, paired, one-tailed. **f**) Fold-changes of primary motor neurons (MNs) survival (relative to each control) after exposure to conditioned media from LPS-stimulated WT and MOK-KO cells (N=3). Student’s t-test performed between shown groups, unpaired, two-tailed. Data in **a**, **c-f** are the mean −/+ S.E.M. from ‘N’ independent experiments, and *P<0.05, **P<0.01 and ***P<0.001. NS: not significant.

To confirm the participation of MOK in the inflammatory response in microglia, CRISPR/Cas9-generated MOK-KO and WT SIM-A9 cells (see **Supplem. Figure 2c,d**) were stimulated with LPS for 5 h and analyzed by qRT-PCR and/or ELISA for a set of key cytokines, namely IL-6, TNFα, IFNβ and IL-1α (**Figure 2c,d, Supplem. Figure 2e,f**), the results clearly demonstrating reduced levels of all these cytokines in LPS-stimulated MOK-KO cells. As expected, this profile of cytokine expression was mirrored by flow cytometry analysis (**Figure 2e, left**). Importantly, we assessed the functional restoration of MOK activity in MOK-KO cells transfected with a FLAG-MOK construct carrying or not a ‘kinase-dead’ (MOK-KD) mutation (25), by IL-6-expression in cells after LPS-stimulation (**Figure 2e, right**). Remarkably, quantification of IL-6^+^ cells in transfected cells (detected with anti-FLAG immunolabelling) demonstrated the rescuing of cytokine expression in MOK-KO cells by recombinant MOK-WT, but not by MOK-KD (**Figure 2e, Supplem. Figure 3**). Put together, these results demonstrate that MOK controls the inflammatory response in microglia and this function is dependent on its kinase activity. To assess the impact in neuronal viability of MOK-regulated microglial responses, we isolated primary motor neurons from mouse embryos for cell culture. We then exposed these cultured neurons to conditioned media from LPS-stimulated WT or MOK-KO cells, and quantified dead cells by using a fluorescence-based assay (**Figure 2f**). As expected, medium from LPS-stimulated WT cells caused a reduction in neuronal cell survival compared to control conditions. Notably, the opposite effect was seen with medium from LPS-stimulated MOK-KO cells, indicating that MOK-mediated microglial responses can be neurotoxic.

We then carried out RNA-Seq analysis by comparing MOK-KO vs. WT SIM-A9 cells, stimulated or not with 1 μg/mL LPS for 5 h (**Figure 3a,b, Supplem. Figure 4**). A relatively large number of protein-coding and non-coding DEGs were identified, showing partial overlap between both cell lines and treatments, and including 730 total DEGs (**Figure 3a, top**) and 424 coding DEGs (**Figure 3a, bottom, 3b**) found when comparing MOK-KO vs. WT cells stimulated with LPS. Gene ontology (GO) clustering analysis revealed significantly enriched biological processes related to microglial activation and innate immunity and, interestingly, IFNα/IFNβ (type-I IFN) regulation and antiviral responses (**Figure 3c,d, Supplem. Figure 5a**). In particular, analysis of coding DEGs that differentially changed for MOK-KO or WT cells upon LPS-stimulation revealed enriched terms related to ‘innate immunity’, ‘positive NF-κB regulation’ and ‘antiviral response’ (**Supplem. Figure 5b**). Finally, IPA prediction between MOK-KO vs. WT cells upon LPS stimulation unleashed a number of activated (CITED2, TRIM24) and inhibited (e.g. IRF7/IRF3, STAT1, BHLHE40) upstream regulators (**Figure 3e**) as well as differentially regulated ‘neuroinflammation’, ‘pyroptosis’ and ‘TRIM1’ canonical pathways, among others (**Supplem. Figure 5c**). The expression of interferon regulatory factor 7 (IRF7), a principal type-I IFN master regulator and also a DEG revealed in our RNA-Seq study, was assessed by qRT-PCR and confirmed to be suppressed in the MOK-KO cells upon LPS stimulation (**Figure 3f**). Collectively, our results demonstrate a role of MOK in microglia activation and inflammatory response mediated by NF-κB and STAT1 pathways and critically involving induction of IRF7/3/type-I IFN and antiviral cellular responses.

**Figure 3.**
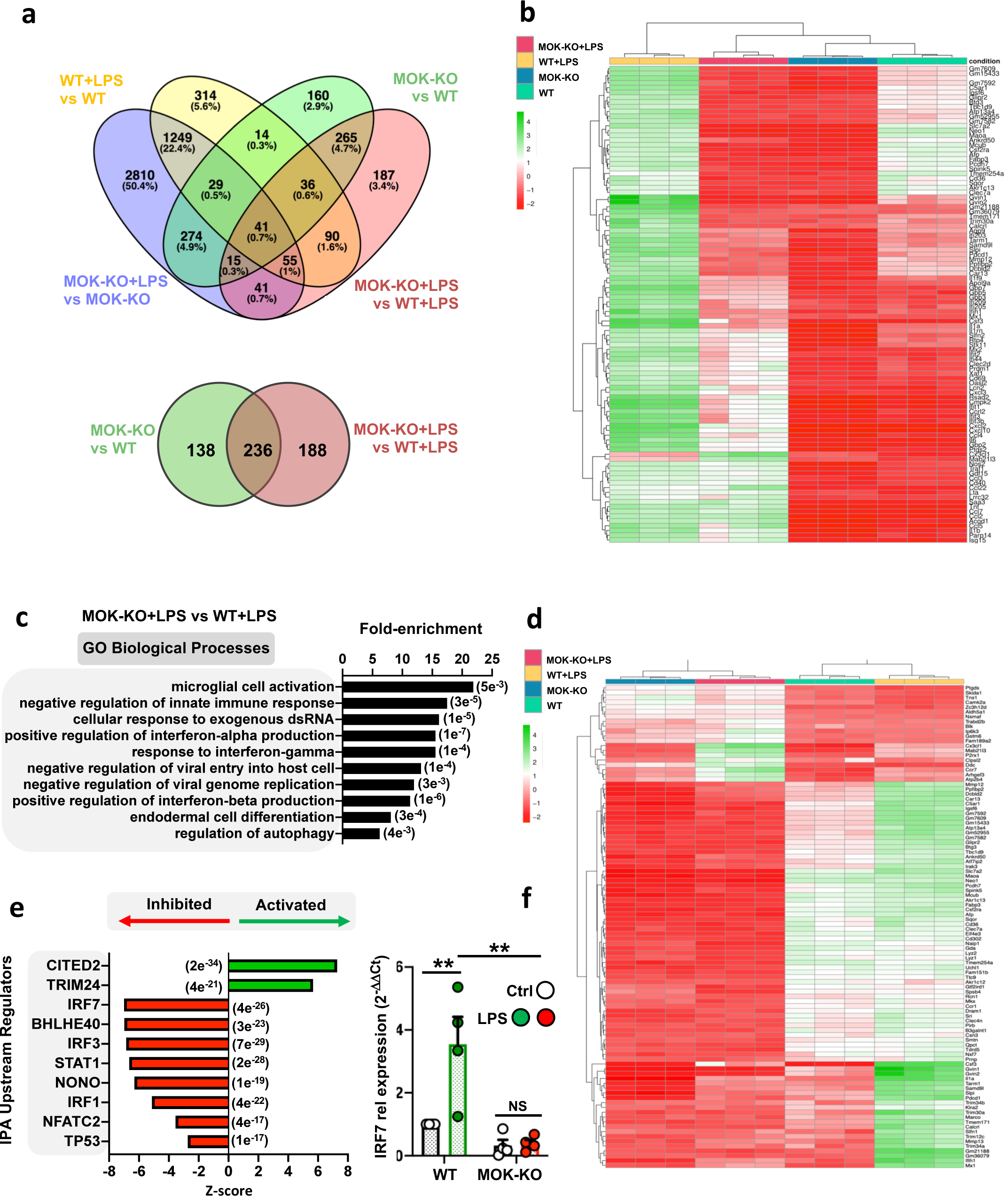
Transcriptomic analysis from RNA-Seq studies of MOK-KO cells and response to LPS stimulation. Data in **a-e** are from 3 independent experiments (N=3). **a**) Venn diagrams depicting the number of total (top) and protein-coding (bottom) DEGs between MOK-KO and WT SIM-A9 cells under basal conditions and/or after 1 μg/mL LPS stimulation for 5 h (pAdj. <0.05; log2 fold-change>2 and <-2); **b**) Heatmap showing unsupervised clustering analysis for the top 100 protein-coding genes based on relative expression levels in the four samples; **c**) GO ‘biological process’ term enrichment analysis of total DEGs for MOK-KO+LPS vs. WT+LPS. Shown are the top hits based on p-value (indicated in parentheses) with at least three up/downregulated genes; **d**) Heatmap depicting clustering analysis for 100 coding DEGs identified for the MOK-KO+LPS vs. WT+LPS comparison (consisting of the top 80% downregulated and top 20% upregulated, genes to maintain proportionality of all significant DEGs found); **e**) Top-10 IPA-predicted upstream expression regulators from coding DEGs comparing MOK-KO vs. WT cells upon LPS stimulation, indicating positive (activated) or negative (inhibited) z-scores. The results are from all genes that were found to be misregulated in the RNA-Seq deseq2 analysis in any comparison with pAdj of <e-5. The P-value of overlap is indicated in parentheses; **f**) Assessment of IRF7 gene expression by qRT-PCR from WT and MOK-KO SIM-A9 cells stimulated with 1 μg/mL LPS for 5 h. Data represent fold-changes relative to unstimulated WT cells (WT Ctrl) (N=4). One-way ANOVA followed by Tukey post-hoc test. Data are the mean −/+ S.E.M. from N independent experiments, and *P<0.05 and **P<0.01.

### MOK mediates immune responses by regulating phospho-Brd4 levels and Brd4 chromatin-binding in microglia

One of the proteins whose phosphorylated state levels we found to be putatively regulated by MOK (**Figure 1c**) corresponds to Brd4 which belongs to the BET bromodomain family, controlling the transcription of genes by binding to acetylated histones in chromatin (26). Brd family members have been linked to cancer and autoimmune disease mechanisms that are still not clear. In particular, Brd4 has been recently involved in innate immune responses, including inflammatory responses (20, 22, 27). To verify that Brd4 is mediating the inflammatory response in our cell line as previously shown in microglia (21, 28), we stimulated SIM-A9 cells in culture with LPS for 5 h after pre-treatment with the Brd4 inhibitor (+)-JQ1 (or (−)-JQ1 negative control) and quantified IL-6 and TNFα secretion levels, confirming that (+)-JQ1 effectively suppresses LPS-induced inflammatory responses in both primary microglia and SIM-A9 cells (**Supplem. Figure 6a,b).**

Next, to validate our screening result and to test whether MOK kinase is indeed regulating phosphorylated Brd4 levels in inflammatory microglia, we quantified Ser^492^ phosphorylated Brd4 (pBrd4) –previously shown to enable Brd4 chromatin-binding functions (29, 30)– in samples from C13-pre-treated SIM-A9 cells stimulated with LPS by Western blot, showing a clear attenuation of the pBrd4 band (ca. 250 kDa) in LPS-stimulated cells (**Supplem. Figure 6c**). Moreover, confocal immunofluorescence analyses with primary microglia showed that, contrary to LPS-stimulated cells displaying higher pBrd4 nuclear levels compared to basal control, C13-pretreatment led to lower pBrd4 nuclear levels following LPS stimulation (**Figure 4a,b).** Importantly, these results were mirrored by LPS-stimulated MOK-KO cells compared to WT cells both by Western blot (**Figures 4c,d, Supplem. Figure 2g**) and confocal immunofluorescence analysis (**Figure 4e,f**), in which a significant reduction of nuclear pBrd4 was seen in MOK-KO compared to WT cells, upon LPS-stimulation. These results clearly demonstrate that MOK kinase regulates nuclear pBrd4 levels in microglia under inflammatory conditions.

**Figure 4.**
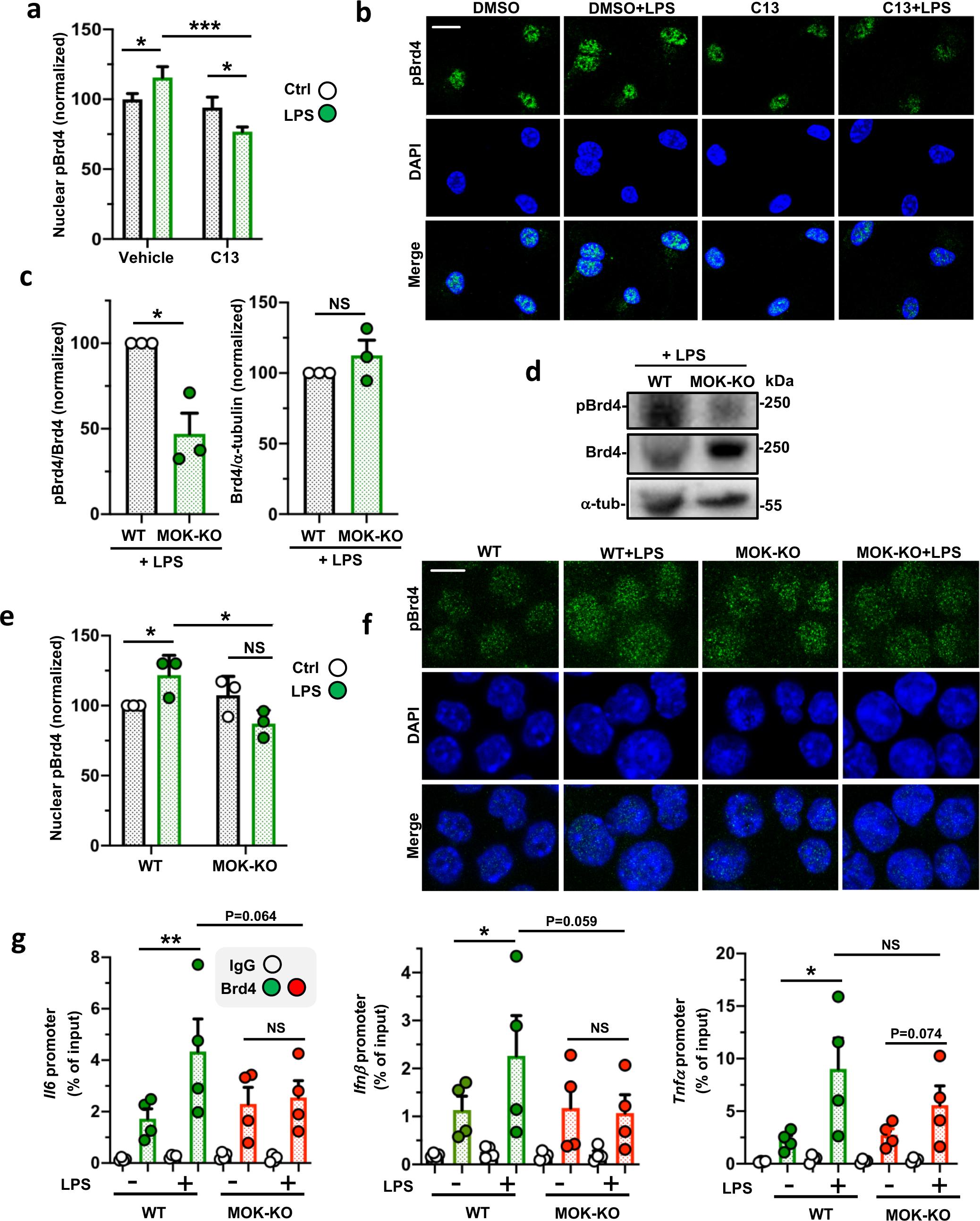
MOK regulates phospho-Ser^492^-Brd4 levels under neuroinflammatory conditions. **a,b)** Quantification of nuclear pBrd4 levels by confocal IF with anti-pBrd4 (green) and DAPI staining of primary microglial cells pre-treated with 10 μM C13 (or DMSO) for 1 h and stimulated or not with 1 μg/mL LPS for 4 h. The data and images shown correspond to one experiment (N=20 analyzed images per condition) and are representative of two independent experiments. Scale bar: 10 μm. **c,d**) Quantification (**c**) and representative image (**d**) of Western blot analyses with anti-pBrd4 and anti-Brd4 (normalized to Brd4 or α-tubulin signal, respectively) in lysates of WT and MOK-KO SIM-A9 cells stimulated with 1 μg/mL LPS for 1 h. Data are from three independent experiments (N=3). Student’s t-test, unpaired, one-tailed. **e,f**) Quantification (**e**) and representative images (**f**) of confocal IF analyses of nuclear pBrd4 levels with anti-pBrd4 (green) and DAPI staining of WT and MOK-KO SIM-A9 cells stimulated or not with 1 μg/mL LPS for 1 h. Signal was enhanced by 50% in all four images in the first row for better visualization. Images are from one out of three independent experiments (N=3). Scale bar: 10 μm. Student’s t-test, unpaired, two-tailed. **g**) ChIP-PCR assessment of Brd4-binding to specific cytokine promoters. Represented data are relative values (percentage of input) of PCR products for *Il6*, *Ifnβ1* and *Tnfα* promoters after ChIP assay with either anti-Brd4 or IgG control antibodies with chromatin isolated from WT or MOK-KO cells treated or not with 1 μg/mL of LPS for 1 hour (N=4). Student’s t-test between shown groups, ratio-paired, one-tailed. Data are the mean −/+ S.E.M. from ‘N’ independent experiments, and *P<0.05, **P<0.01 and ***P<0.001. NS: not significant.

To assess a possible role of MOK in Brd4 binding to key cytokine gene promoters previously linked to Brd4 recruitment (31–33), we carried out ChIP-qPCR assays from SIM-A9 cells after 1h-LPS stimulation (**Figure 4g**). Remarkably, we found that whereas WT cells showed incremented Brd4 binding to *Il6*, *Ifnβ1* and *Tnfα* gene promoters in response to LPS compared to unstimulated cells, such increase was abrogated in MOK-KO cells for *Il6*, *Ifnβ1* and to a lesser extent, *Tnfα*, promoters. These results demonstrate a role of MOK in positive regulation of Brd4 binding to proinflammatory and type-I IFN gene promoters under inflammatory conditions. Of note, pre-treatment with the Brd4-inhibitor (+)-JQ1 resulted in significant reduction of the ChIP-qPCR signal increments in all cases, confirming the specificity of Brd4 binding assessment in our assays (**Supplem. Figure 6d**). Collectively, these results reveal a central role for MOK in Brd4-mediated regulation of cytokine expression upon inflammatory stimulation. On the other hand, the lack of a significant decrease in Brd4 binding to TNFα promoters in LPS-stimulated MOK-KO compared to WT cells suggests that Brd4-independent, MOK-mediated mechanisms are also contributing to the upregulation of certain cytokines during microglial responses.

### MOK is altered in spinal cord from ALS patients and mouse model, and C13-administration to SOD1^G93A^ mice modifies disease course

Given that MOK was found to be engaged in microglia under TDP-43 aggregation and based on our results herein showing a role of MOK in the inflammatory response, we wondered whether MOK would be altered in ALS individuals. To address this question, we first assessed phospho-Ser^159^/Tyr^161^ MOK (bpMOK), considered to correspond to the fully active form of the kinase (14). Western blot analysis using murine spinal cord tissue homogenates showed a band of ca. 65 kDa, in agreement with previously reported MW for CNS tissue (34) (**Supplem. Figure 7a**). Next, we carried out immunohistochemistry assays to detect bpMOK in spinal cord tissue from a TDP-43-based mouse model (Tg^TDP43^) (35) at the late-stage and from sporadic ALS patients. Considering the significant differences in cell size measured between WT and Tg^TDP43^ samples (**Supplem. Figure 7b**), we normalized the quantitated bpMOK levels in these samples to the cell area. Remarkably, the results revealed significantly higher levels of bpMOK in the mouse model compared to age-matched mice (**Figure 5a**). Furthermore, assessment of bpMOK cellular signal in tissue from human subjects also showed increased bpMOK levels in ALS samples compared to age-matched controls (**Figure 5b**).

**Figure 5.**
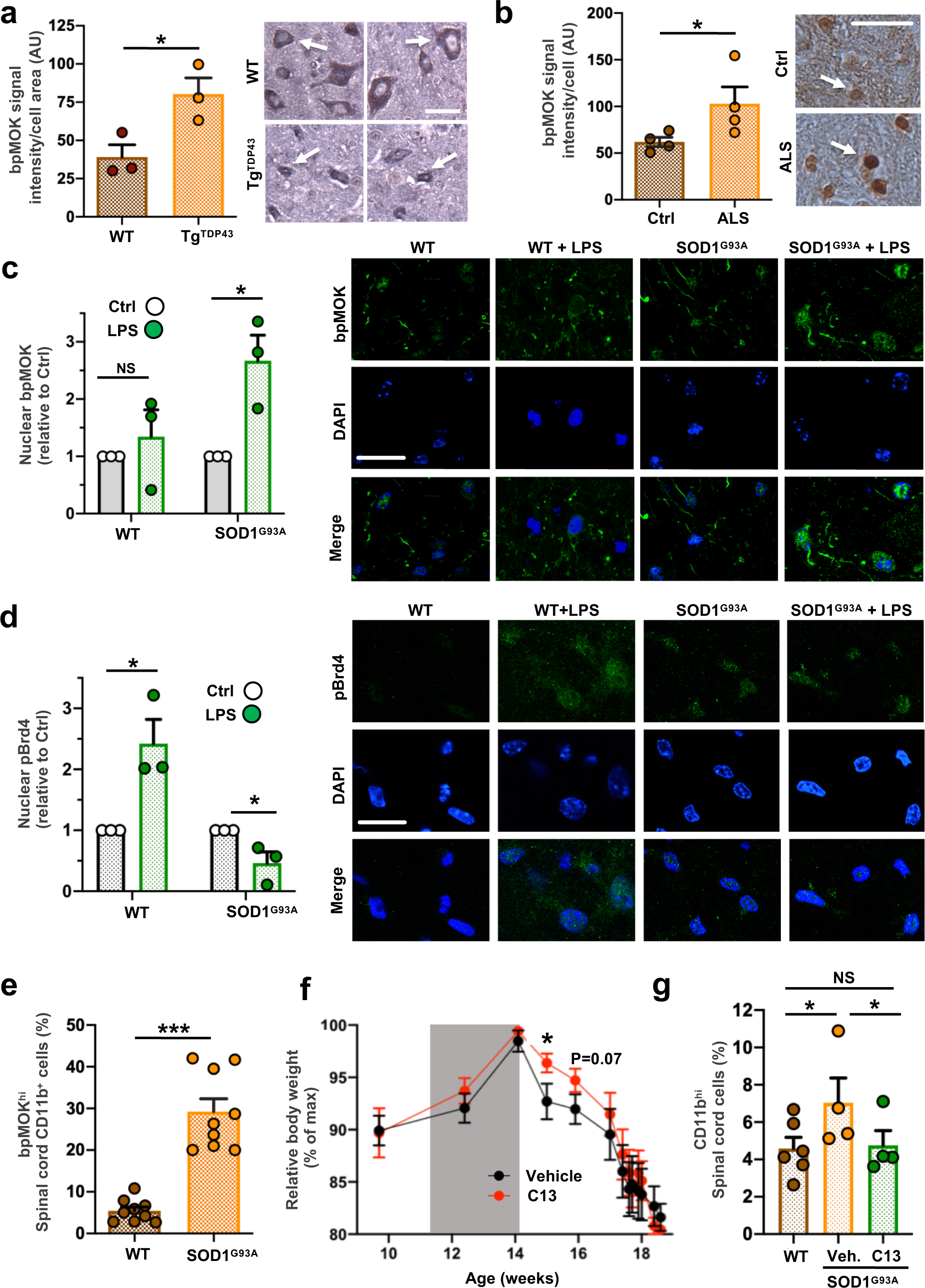
MOK is altered in CNS cells in the context of ALS. Quantification of bpMOK levels by IHC analysis in spinal cord tissue samples (left) and representative IHC images (right) for spinal tissue samples from **a**) Tg^TDP43^ and WT mice (late stage; N=3) and **b**) sporadic ALS patients and age-matched control subjects (N=4). Student’s t-test, unpaired, one-tailed. Scale bar: 25 μm. White arrow indicates one analyzed cell as an example. **c, d**) Left: quantification of nuclear bpMOK (**c**) or pBrd4 (**d**) levels (green) and DAPI staining by confocal IF of spinal cord organotypic cultures, stimulated or not with 1 μg/ml LPS overnight, from WT and SOD1^G93A^ mice (5-weeks old, N=3). Represented data are the fold-change of nuclear bpMOK signal upon LPS stimulation normalized to the corresponding unstimulated controls. Student’s t-test, unpaired, two-tailed. Right: representative images of both IF assays corresponding to each condition. Scale bar: 25 μm. **e**) Flow cytometry analysis of the bpMOK^hi^-expressing population in CD11b^+^ cells isolated from early-onset SOD1^G93A^ or WT mouse spinal cords (14-weeks old; N=9). Student’s t-test, unpaired, two-tailed. **f**) Time-course body weight monitorization of SOD1^G93A^ mice injected (i.p.) with C13 compound (20 μg per dose) or vehicle (N=6) starting at 11 weeks of age, every other day along three weeks (grey window). All body weight values are normalized to each mouse’s maximum reached weight (% of maximum). One-way ANOVA (mixed-results model) followed by Tukey post-hoc test. **g**) Flow cytometry analysis of the CD11b^hi^-expressing population in live cells isolated from spinal cord from WT or C13- or vehicle-administered SOD1^G93A^ mice (14-weeks old; N=6 for WT, N=4 for C13/vehicle-treated SOD1^G93A^ mice). The CD11b^hi^ signal threshold corresponded to the top 5% (on average) for the WT mice. Student’s t-test, paired, one-tailed. Data are the mean −/+ S.E.M. from ‘N’ independent experiments, and *P<0.05, **P<0.01 and ***P<0.001. NS: not significant.

To get an insight into a possible engagement of MOK in a pathophysiological context of ALS neuroinflammation, we chose the well-established and neuroinflammation-recapitulating SOD1^G93A^ animal model (24, 36–38), which has additionally been found to display TDP-43 functional abnormalities (39–43), and investigated the relative changes in bpMOK levels induced by LPS-stimulation in spinal cord organotypic cultures from pre-symptomatic (5-weeks old) SOD1^G93A^ mice vs. WT controls, by IF confocal analysis (**Figure 5c**). As opposed to WT mice, in which no significant changes were observed when stimulated with LPS, a significant increase in nuclear bpMOK was seen in LPS-stimulated spinal cultures from ALS mice. Remarkably, by performing an analogous study with pBrd4 immunolabeling (**Figure 5d**) we observed that, whereas LPS stimulation resulted in upregulated nuclear pBrd4 levels in spinal cord tissue from healthy mice, no upregulation -but rather, a reduction-in pBrd4 was induced for ALS animals. These results suggest that the MOK-pBrd4 axis activated in the inflammatory response is dysregulated in ALS.

To determine whether the levels of functional MOK kinase could be specifically altered in microglia in the context of ALS, we isolated microglial cells from spinal cord of WT or SOD1^G93A^ mice at post-onset (14 weeks-old) and quantified the cells displaying high levels of bpMOK among the cell population expressing the microglial surface marker CD11b, by flow cytometry (**Figure 5e, Supplem. Figure 7c,d**). Remarkably, bpMOK^high^ microglial cells were significantly more abundant in cells isolated from SOD1^G93A^ mice compared to controls, showing ca. 5-fold increase in the ALS model.

Finally, to investigate a possible role of MOK in the pathophysiology of ALS *in vivo* and its potential as a therapeutic target, we peripherally administered the MOK inhibitor C13, or vehicle, to SOD1^G93A^ mice by intraperitoneal injection every other day along three weeks, starting at the pre-onset stage (11 weeks of age). Assessment of motor performance by Rotarod test did not show any significant differences between both groups (**Supplem. Figure 7e**). Notably, however, C13-treatment resulted in protection during a time window at the post-onset stage and reaching statistical significance one week after the end of treatment, according to relative body weight assessment, another standard parameter used to monitor disease course in this ALS model (**Figure 5f**). Finally, to investigate whether the protective effects of C13 administration to SOD1^G93A^ mice might reflect suppressed microgliosis, we analysed the content of CD11b^high^ cells in spinal cord from treated or non-treated mice past week 14, i.e. just after the end of treatment (**Figure 5g**). As expected, flow cytometry analysis showed increased CD11b^high^ cell content in SOD1^G93A^ compared to WT mice, indicating increased microglia activation in ALS spinal cord (44). Remarkably, the analysis also revealed suppressed levels of CD11b^high^ in SOD1^G93A^ mice spinal cord as a result of C13 administration, suggesting that MOK is indeed involved in microglial activation and neuroinflammation in the context of ALS. Of note, analysis of brain motor cortex samples from these mice revealed overtly lower pBrd4 levels by IF in C13-treated compared to vehicle-treated, ALS mice (**Supplem. Figure 7f**). All together, these results demonstrate that MOK kinase and its newly unleashed regulated pathways are altered in ALS spinal cord cells, including microglia, and they indicate that MOK is involved in the pathophysiology of ALS.

## Discussion

Previously, we identified the Ser/Thr kinase MOK as a protein that strongly interacts with abnormal cytoplasmic TDP-43 aggregates in microglial cells (15). Until now, no regulated signalling pathway or downstream molecular target has been identified for MOK, and its potential function in immune cells has not been explored. In this study we obtained a differential profile of phosphorylated protein candidates under TDP-43 aggregation conditions for C13 pre-treated cells, suggesting that MOK is regulating downstream phosphorylation events in microglia under TDP-43 aggregation, including the epigenetic reader Brd4. Furthermore, pre-treatment of microglial cells with MOK inhibitor C13 revealed a number of genes whose early expression upon exposure to TDP-43 aggregates would be regulated by MOK, including Dtx2 (regulator of Notch signalling), Ppp5c (Ser/Thr phosphatase involved in various signalling pathways) and Sap30I (transcription repressor via interaction with histone-deacetylase complexes/HDACs). Together, these data demonstrate that MOK kinase is functionally mobilized in microglial cells under TDP-43 aggregation conditions, which suggests a role of this kinase in the pathogenic mechanisms triggered by TDP-43 aggregation processes which include eliciting proinflammatory responses.

Indeed, by using a uniquely specific inhibitor of MOK kinase (18) and MOK-deleted cells, we showed that MOK kinase controls the inflammatory response induced by the general stimulator LPS in microglia. Moreover, by exogenous expression of recombinant WT or kinase-dead MOK protein in MOK-KO cells, we demonstrated that MOK regulates proinflammatory cytokine expression in a kinase dependent manner. By supporting IL-6, TNFα and IL-1α/β along with the activation of the p65/RelA NF-κB pathway, MOK regulates the canonical inflammatory response, while also supporting IFNβ secretion, which has been lately related to neuroinflammatory processes and disease progression in ALS and other neurodegenerative disorders (45–47). Integrated transcriptomics analysis of enriched biological processes and KEGG pathways indicated that MOK-dependent control of the inflammatory response in microglia is mediated by regulation of microglial activation, the ‘innate immune response’, IFNα/β (type-I IFN) production, and anti-viral responses. IPA prediction of upstream regulators supports that MOK mediates the ‘neuroinflammation signalling pathway’ in LPS-stimulated microglia by stirring the inhibition of CITED2 and TRIM24 —a critical negative regulator of inflammatory gene programs in myeloid cells (48–50) and a suppressor of M2 macrophage polarization (51), respectively. In the case of C13-treated microglia, the top predicted MOK-inhibited regulators were CEBPB, NFE2L2 and STAT3. Consistent with an NF-κB-mediated anti-inflammatory effect of C13 in LPS-stimulated microglia, CEBPB has been recently found to be driven by the NF-kB pathway in dendritic cells in the context of inflammation (52) and has been reported to mediate macrophage M2-phenotype polarization (53, 54) and microglial Aβ phagocytosis (55). Recently, STAT3 was shown to act coordinately with CEBPB (56) to activate the NFKBIZ/IκB-ζ protein involved in the regulation of inflammatory responses (57), whereas NFLE2L2/Nrf2 is known to elicit anti-inflammatory responses in microglia and other immune cells (58). In addition, as revealed in both transcriptional studies with MOK-KO cells and C13-treated primary microglia, MOK favours the activation of STAT1 and IRF7/3 upstream regulators, which are well known IL-6- and type-I IFN-promoting transcription factors, respectively (59). Indeed, the suppression of IFNβ secretion and the type-I IFN transcriptional signature in MOK-deficient inflammatory microglia indicates that MOK regulates the type-I IFN response, which has been lately related to neuroinflammatory processes in a variety of neurodegenerative situations, including ALS, Alzheimer’s disease and other tauopathies, Parkinson’s disease, prion-disease and traumatic brain injury (60–67). BHLHE40, another predicted MOK-activated regulator, has been reported to act as a central transcription factor for pro-inflammatory gene expression in macrophages and other immune cells (68, 69). Moreover, it has been recently found to be involved in the regulation of the MGnD microglial neurodegeneration-associated (DAM) expression signature (38).

Remarkably, we identified the first protein whose phosphorylation state levels are physiologically regulated by MOK, the epigenetic reader Brd4. Brd4 was previously demonstrated to be phosphorylated at Ser^492^ by casein kinase II/CK2 (30). Therefore, our results indicate that MOK could possibly be acting on pBrd4 levels directly via its Ser/Thr kinase activity or indirectly, possibly by regulating CK2 activity.

Notably, Brd4 has been lately shown to play crucial roles in innate immune responses by activating transcription of a number of immune genes (26, 70) and BET proteins dysfunction has been lately involved in various inflammation-related disorders, including cancer, neurodegenerative disease and CNS injury (26, 27). In this study, we showed that MOK supports nuclear levels of the critical phospho-Ser^492^-Brd4 (pBrd4) state and, furthermore, we demonstrated that MOK controls Brd4 binding to cytokine gene promoters in proinflammatory microglia. This unleashed regulation of Brd4 promoter-binding and concomitant cytokine-expression by MOK upon acute or ‘non-pathological’ inflammatory stimulation could be of utmost relevance as evidence toward an impact of epigenetic factors in ALS and other neurodegenerative disorders has already started to emerge (71, 72). Intriguingly, the MOK-dependent regulation of TNFα secretion we found not to rely on Brd4’s promoter-binding function may still be Brd4-dependent, as Brd4 was reported to control the NF-κB-inhibitor IkBα at the translation stage, in inflammatory macrophage responses (20).

Finally, we showed that the levels of (Ser^159^Tyr^161^)-phospho-MOK (bpMOK), either constitutive or induced by proinflammatory stimulus, are significantly altered in the spinal cord of ALS patients or animal models, indicating that MOK’s physiological functions are dysregulated in the context of ALS. In particular, we reveal a massive increase of bpMOK levels in microglia of SOD1^G93A^ mice. Moreover, we observed that inflammatory stimulation of spinal cord samples from pre-onset SOD1^G93A^ mice, as opposed to WT mice, fails to upregulate nuclear pBrd4 levels, indicating dysfunctional or saturated MOK-Brd4 signalling in the ALS model. Remarkably, we observed that peripheral administration of C13 MOK-inhibitor to pre-onset ALS mice results in abrogated pBrd4 levels and reduced CD11b^high^-microglia levels in CNS, while significant protection from disease progression was seen along a short period after treatment was discontinued. These results strongly indicate that MOK is involved in ALS pathophysiology and suggests that targeting MOK with a specific inhibitor in a continued manner could be beneficial throughout the disease course. The broad relevance of our findings is further evidenced by two recent studies revealing associations between MOK gene expression levels in peripheral leukocytes with blood pressure (73) and with type-1 diabetes (74), which supports a possible role for MOK in immune dysregulation in the context of other pathological scenarios. Indeed, protein kinases, including immune-related kinases, have been shown to play crucial roles in the physiopathology of ALS and other neurodegenerative proteinopathies and have recently emerged as attractive therapeutic targets in various neurodegenerative disorders (75, 76). Therefore, the identification of MOK-regulated proteins and downstream signalling pathways in immune cells, particularly in the context of neuroinflammation, could set the bases for therapeutic targeting of key pathogenic pathways in ALS and other inflammation-related disorders.

Overall, we unleash a role of MOK kinase -the first related to immunity or the CNS-in controlling the inflammatory response in microglia via Brd4-dependent and independent mechanisms, and report that MOK microglial levels and its immune-related functions are dysregulated in ALS, with strong implications in disease pathophysiology. This constitutes a novel signalling pathway in neuroinflammation and discloses new potential intervention targets in ALS and other neurodegenerative and inflammatory diseases.

## Methods

### General reagents and DNA constructs

Lipopolysaccharide (LPS) from *Escherichia coli* (L3137, Sigma-Aldrich), (−)-JQ1 (SML1525, Sigma-Aldrich), (+)-JQ1 (SML1524, Sigma-Aldrich), polymixin B solution (81271, Merck Life Science S.L.) and dimethyl sulfoxide (DMSO, D2650, Sigma-Aldrich) were used. Compound AG2P145D/Comp13 (C13) was synthesized and characterized as previously reported (18). Plasmids encoding FLAG-tagged MOK (murine, wild type/WT and kinase-dead/KD) were described previously (25).

### Human samples

Patients spinal cord samples were obtained after voluntary donation to the Brain Bank of the Region of Murcia (BCRM) by patients diagnosed as sporadic ALS. Control spinal samples from subjects with no history of neurological diseases were donated by the BCRM. The sample collection at BCRM fulfils the ethical standards of our institutions: IMIB-Arrixaca and ISCIII National Biobank Network Review Boards approved the protocol (anonymization/custody/conservation in control samples and custody/conservation in patient’s samples), which was required for Clinical Trial approval (NCT00124539, EudraCT 2006-00309612), as well as the 1964 Helsinki declaration and its later amendments or comparable ethical standards. Prior to the spinal cord extraction, *in situ* exploration was performed to detect possible anatomical malformations. Transverse serial segments of 1.0 cm were embedded in paraffin, cut at 7 μm and mounted in 10 parallel series for IHC immunolabeling.

### MOK-knockout cells

MOK-knockout (MOK-KO) SIM-A9 cell clones (H14 and A7) were generated by CRISPR/Cas9 technology (Synthego, Redwood City, CA, USA). Quality control of MOK-KO clones was done by Synthego. The expected mutations were confirmed by Sanger DNA sequencing (Genomics Unit, CNIO, Madrid) and clones were validated by MOK-specific qRT-PCR and Western blot by us, as shown in Supplem. Figure 2c,d.

### Animal models

For IHC studies, spinal cord tissue slices (lumbar, sacral) were from late-stage Prp-TDP43^A315T^ mice (Tg^TDP43^), an ALS/FTD model described previously (35), and wild-type, age-matched controls. For all the remaining studies, transgenic B6SJL-TgN(SOD1-G93A) 1Gur/J (Jackson Laboratory, USA) male mice carrying the human SOD1 gene with the G93A point mutation (SOD1^G93A^) (77) and wild-type counterpart mice were used. Housing, breeding, and animal welfare approvals were as detailed in SI Appendix.

### Administration of C13 to SOD1^G93A^ mice and phenotype monitorization

Each mouse was injected intraperitoneally every other day with a total of 100 μl solution consisting of either 20 μg of C13 compound (18) in PBS or vehicle control containing 2% DMSO, for a total of three weeks starting at 11 weeks of age. Mice were weighed and tested in rotarod twice a week. At the end of treatment, a number of mice from each group were sacrificed and their spinal cord and brain were removed for microglial cell isolation followed by CD11b assessment by flow cytometry and for pBrd4 analysis by IF, respectively (as described in Supplem. Methods). The remaining mice continued to be assessed by rotarod and weighed regularly until they reached 80% of their maximum weight, at which point they were sacrificed.

### Cell culture

Mixed glial cultures were prepared from cerebral cortices of 1 day- to 3 day-old C57BL/6 male mice (University of Seville Animal Core Facility), and microglia were isolated by mild trypsinization as previously described (15). After microglia isolation, cells were detached with accutase (A11105, Thermo Fisher Scientific) and seeded in 24-well plates at a density of 100,000 cells/mL. Attached microglia were allowed to recover in conditioned medium (half new culture medium, half mixed microglia-used culture medium) for 48 h before being subjected to different treatments. SIM-A9 microglial cell line (CRL-3265) was acquired from ATCC. SIM-A9 WT cells and MOK-KO clones were maintained in DMEM-F12 (Sigma-Aldrich) that was supplemented with heat-inactivated 10% FBS and 5% HS (Gibco, Billings, MT, USA), 2 mM L-glutamine (Thermo Fisher Scientific), 100 U/ml penicillin (Thermo Fisher Scientific) and 100 μg/ml streptomycin (Thermo Fisher Scientific).

### Treatment of cell cultures with LPS

Stimulation of primary or SIM-A9 microglial cultures with LPS was performed by adding one tenth volume of medium alone or containing either LPS (1 μg/mL final) and incubated for either 1, 5 or 16 h, as indicated. Pre-treatment with 10 μM C13 or 10 μM (−)-JQ1/(+)-JQ1 was done in the same manner by incubating cells for 1 h prior to cell treatment. In these cases, control cultures received equal volume of DMSO in medium (vehicle). After incubation at 37°C, supernatants were harvested, clarified by mild centrifugation and stored at −20°C, and cells were either used for different purposes or frozen and stored at −20°C. For cell treatment with TDP-43 aggregates for SLAM-Seq analysis, see below.

### Preparation of TDP-43 aggregates and treatment of cell cultures

TDP-43 and sham aggregates were obtained by purifying the inclusion body fraction after overexpression of His_6_-human TDP-43 fusion protein (by using a plasmid construct kindly provided by Yoshiaki Furukawa’s laboratory, Keio University, Yokohama, Japan), or empty vector, in *E. coli* BL21(DE3) cells as previously described (15, 78, 79). General characterization of the *in vitro*-prepared TDP-43 aggregates was done as before (15). SIM-A9 or primary microglial cells in culture were pre-treated with 10 μM final C13 or DMSO for 1 h and then to 5 μg/mL TDP-43 or sham aggregates in medium containing polymyxin B (10 μg/mL) 5 h or overnight. Culture supernatants were harvested for IL-1β and IL-18 quantification by ELISA and Western blot, respectively, and cells were lysed for GAPDH assessment by Western blot.

### Organotypic culture preparation and treatment

Organotypic cultures were prepared as previously described(15) from lumbar spinal cords of 5-week-old male SOD1^G93A^ or wild-type (WT) mice. Cultures were maintained at 37°C in a 5% CO_2_/95% air humidified environment. Cultures were left to stabilize for 4 days, then medium was changed every 3 days. After 1 week, treatments were performed by adding one tenth volume of fresh medium containing LPS (1 μg/mL final) or medium alone. After overnight incubation, supernatants were harvested and significantly upregulated IL-6 cytokine levels were confirmed by ELISA in all LPS-treated cultures as a stimulation check-up. In addition, organotypic cultures were processed for immunofluorescence analyses (see below).

### Acute microglial cell isolation from adult mice

Microglial cells from adult mice were acutely isolated from spinal cord tissues of 14 weeks-old mSOD1^G93A^ Tg mice and their WT littermates. A detailed description can be found in SI Appendix.

### Immunolabeling of acutely isolated microglial cells and flow cytometry analysis

A detailed description of immunolabeling of CD11b and/or bpMOK can be found in SI Appendix.

### Immunohistochemistry (IHC) and IHC analyses

IHC assays and quantification were double blinded. Fixed spinal cord segments from Tg^TDP-43^ and control mice and controls (lumbar region) or human sporadic ALS patients were used for immunolabeling and quantification of bpMOK. A detailed description can be found in SI Appendix.

### Chromatin immunoprecipitation (ChIP) and ChIP-qPCR

SIM-A9 cell cultures were crosslinked with 1% (v/v) formaldehyde in crosslinking buffer (50 mM Hepes pH 8.0, 0.1 M NaCl, 0.1 mM EDTA pH 8.0, 0.5 mM EGTA pH 8.0). After 10 min of incubation at 37°C, the crosslinking reaction was stopped with 125 mM glycine for 5 min at room temperature. After washing twice with cold PBS, cells were collected by scraping in PBS supplemented with Protease inhibitor cocktail (PIC, Sigma-Aldrich, Cat. 11836170001) and PMSF (ITW Reagents, Cat. A0999). Cells were lysed by consecutively resuspending the pellet in two different lysis buffers (5 mM PIPES pH 8.0, 85 mM KCl, 0.5% IGEPAL and PIC, and 1% SDS, 10 mM EDTA pH 8.0, 50 mM Tris-HCl pH 8.1 and PIC) with a centrifugation step in between. Lysates were sonicated with Digital Sonifier 450 (BRANSON) to obtain chromatin fragments of 200-500 bp. Samples were incubated with anti-BRD4 antibody (Bethyl) and retrieved with Protein G-Dynabeads (Thermo Fisher, Cat. 1004D). After the cross-linking was reversed, chromatin fragments were treated with proteinase K. DNA fragments were purified using a ChIP DNA Clean & Concentrator Kit (Zymo Research, Cat. D5205). qPCR was performed as described above. In order to analyze the qPCR data of immunoprecipitated samples, normalization to the input DNA was performed. Primers pairs (Sigma-Aldrich) specific for *Il6*, *Tnfα* and *Ifnβ1* gene promoters were designed to define an amplicon of 100-170 bp (Table S3).

### Immunofluorescence (IF) of cultured cells

Immunolabeling was done for pBrd4, followed by confocal fluorescence analysis. A detailed description can be found in SI Appendix.

### Immunofluorescence (IF) of spinal organotypic culture and brain slices

Immunolabeling was done for bpMOK or pBrd4 in samples from WT or SOD1^G93A^ mice. A detailed description can be found in SI Appendix.

### Cytokines determination by ELISA

A detailed description of IL-6, TNF-α, IFN-β and IL-1α can be found in SI Appendix.

### Transfection of microglial cell cultures

For the overexpression of FLAG-tagged WT-MOK and KD-MOK constructs, Glial Mag transfection reagent (OzBiosciences, Cat. KGL0250) was used. A detailed description can be found in SI Appendix.

### Immunolabelling and Flow Cytometry of transfected cells

A detailed description of flow cytometry analysis of IL-6 immunolabelled-SIM-A9 cells can be found in SI Appendix.

### Preparation of cell lysates and Western Blot assays

A detailed description can be found in SI Appendix.

### Cell lysate preparation for anti-phospho-Ser/Thr immunoprecipitation

A detailed description can be found in SI Appendix.

### LC-MS/MS and protein identification

A detailed description of processing and analysis of IP protein eluates can be found in SI Appendix.

### RNA isolation and cDNA synthesis for qRT-PCR, qRT-PCR and RNA-Seq analysis

A detailed description of these procedures can be found in SI Appendix.

### RNA-Seq and mRNA-Seq data processing

A detailed description can be found in SI Appendix.

### Gene set enrichment analyses. Gene ontology (GO), pathway enrichment and ingenuity pathway analysis (IPA)

For each comparison group of SIM-A9 cells (WT, WT+LPS, MOK-KO, MOK-KO+LPS), gene ontology and pathway enrichment analyses were performed using the R-package PathfindR (v1.6.2). All differentially expressed genes (DEGs) obtained after applying the filter on Log2 Fold Change values >2 or <-2 along with P value <0.05 were used as input to perform enrichment for the following categories: GO Biological Process, GO Cellular Component, GO Molecular Function and KEGG Pathway with analyses performed by Bencos Research Solutions Pvt. Ltd. (Thane, Maharashtra 400615, IN). Ingenuity Pathway Analysis was applied on DEGs as defined above.

### S4U-labeling and SLAM-Seq analysis

A detailed description can be found in SI Appendix.

### Statistical analyses

Data are presented as mean ± SEM of N independent experiments, unless indicated otherwise. Statistical analyses were performed using either One-way ANOVA followed by Tukey’s post-hoc test or Student’s t-test in the GraphPad Prism software, two-/one-tailed as indicated in the corresponding figure legend. P values less than or equal to 0.05 were considered statistically significant. Statistical analyses were performed using the GraphPad Prism software v.8 (GraphPad Software, La Jolla, USA).

## Acknowledgments and funding sources

We are extremely grateful to Prof. Kevan Shokat and Dr. Flora Rutaganira, University of California, San Francisco (UCSF, San Francisco, USA) for kindly supplying us with C13 compound. We thank patient associations (Saca la lengua a la ELA, Juntos contra la ELA and Reto Todos Unidos/Miquel Valls Fundació Catalana D’ELA) for supporting PAIDI Research Group (CTS-0160) global mission. Financial support to C.R. was provided by the Spanish Ministry of Economy (RTI2018-098432-B-I00), Fundación Ramón Areces (CIVP19A5938), the Andalusian Regional Government-FEDER (US-1265227), I+D PAIDI (PY20-01097). Funding to D.P. was given by PAIDI research group (CTS-0160) and Regional Ministry of Health (PI-0232-2022). V.C-G. is supported by the Institute of Health Carlos III, Spain, co-funded by the European Social Fund (CP19/00046). S.M. was supported by Generalitat Valenciana (GVA: Prometeo/2018/041 and CIPROM/2021/018); MINECO/AEI/ERDF, EU, the Spanish Ministry of Economy, Industry and Competitiveness, the Spanish State Research Agency co-funded by ERDF (PID2020-118171RB-100); and Instituto de Salud Carlos III (RD16/001/0010, co-funded by ERDF).

