## Supplemental Information for "MAPK/MAK/MRK overlapping kinase (MOK) controls microglial inflammatory/type-I IFN responses via Brd4 and is involved in ALS"

**This PDF file includes:**

Figures S1 to S7
Supporting Materials and Methods
Tables S1 to S3

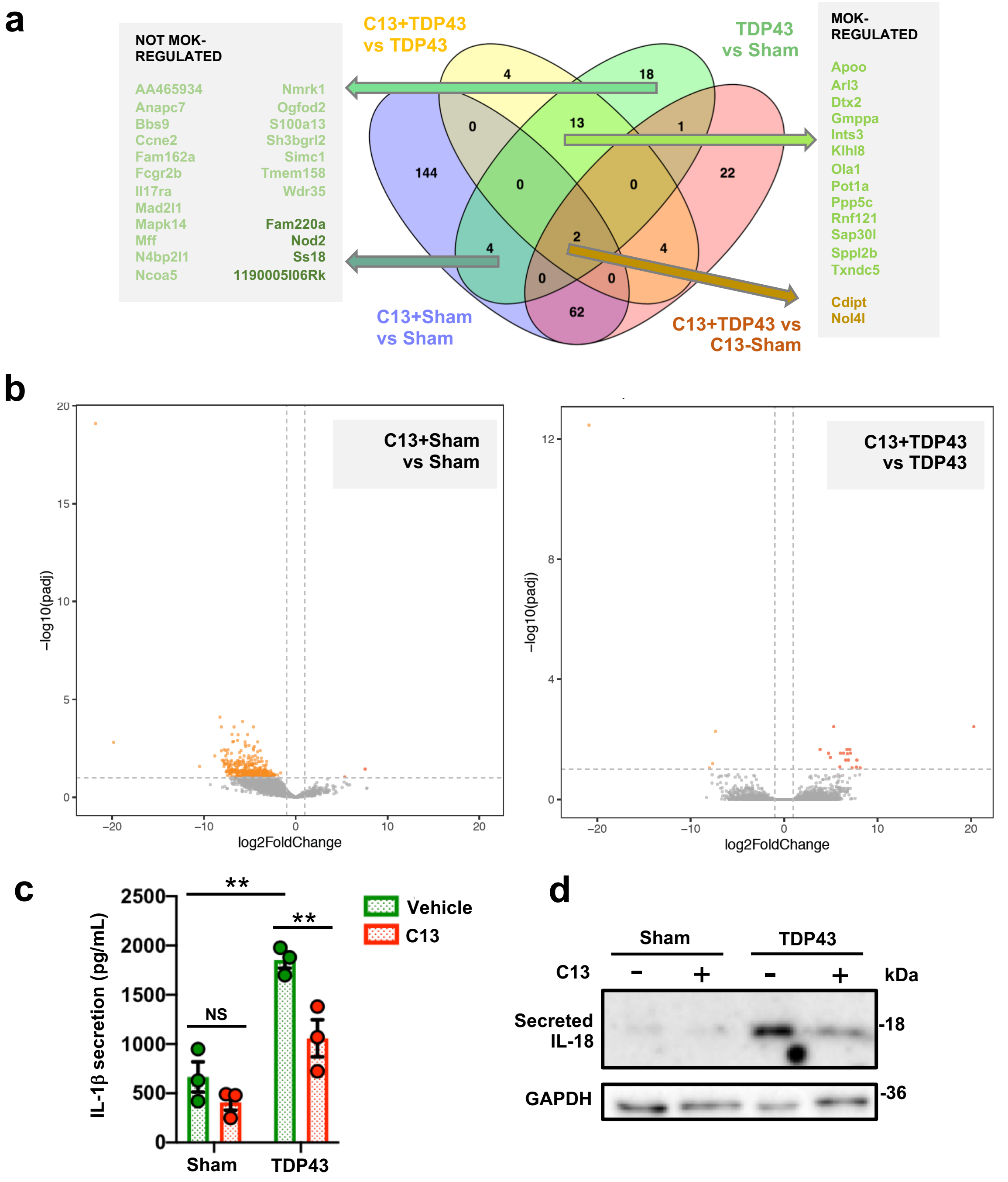


**Supplem. Figure 1: a)** Venn diagrams (created with Venny's [80]) showing the number of DEGs from analyzed SLAM-Seq data obtained with SIM-A9 cells exposed to 5 μg/mL TDP-43 aggregates (TDP43) or sham aggregates (Sham) for 4 h, pre-treated for 1 h with 10 μM C13 or DMSO (vehicle). Results are from three independent experiments (N=3). Indicated are the number of DEGs for 'C13+Sham vs. Sham' and 'C13+TDP43 vs. C13+Sham' comparisons (pAdj.<0.05) and for the 'TDP43 vs. Sham' and 'C13+TDP43 vs. TDP43' comparisons (pAdj.<0.1). Lists of DEGs for 'not MOK-regulated' (left) and 'MOK-regulated' DEGs within those genes that change upon exposure to TDP-43 aggregates. **b)** Volcano plots representing the identified DEGs (with pAdj.<0.05) for 'C13-Sham vs. Sham' (left) and 'C13+TDP43 vs. TDP43' (right). **c)** Determination of IL-1β by ELISA (N=3) and IL-18 by Western blot (representative image of two experiments) from primary microglial cells stimulated with 5 μg/mL TDP-43 aggregates (TDP43) or sham aggregates (Sham) overnight, pre-treated for 1 h with 10 μM C13 or DMSO (vehicle). Data are mean -/+ S.E.M. Student's t-test between shown groups, unpaired, one-tailed. **P<0.01, NS: not significant.

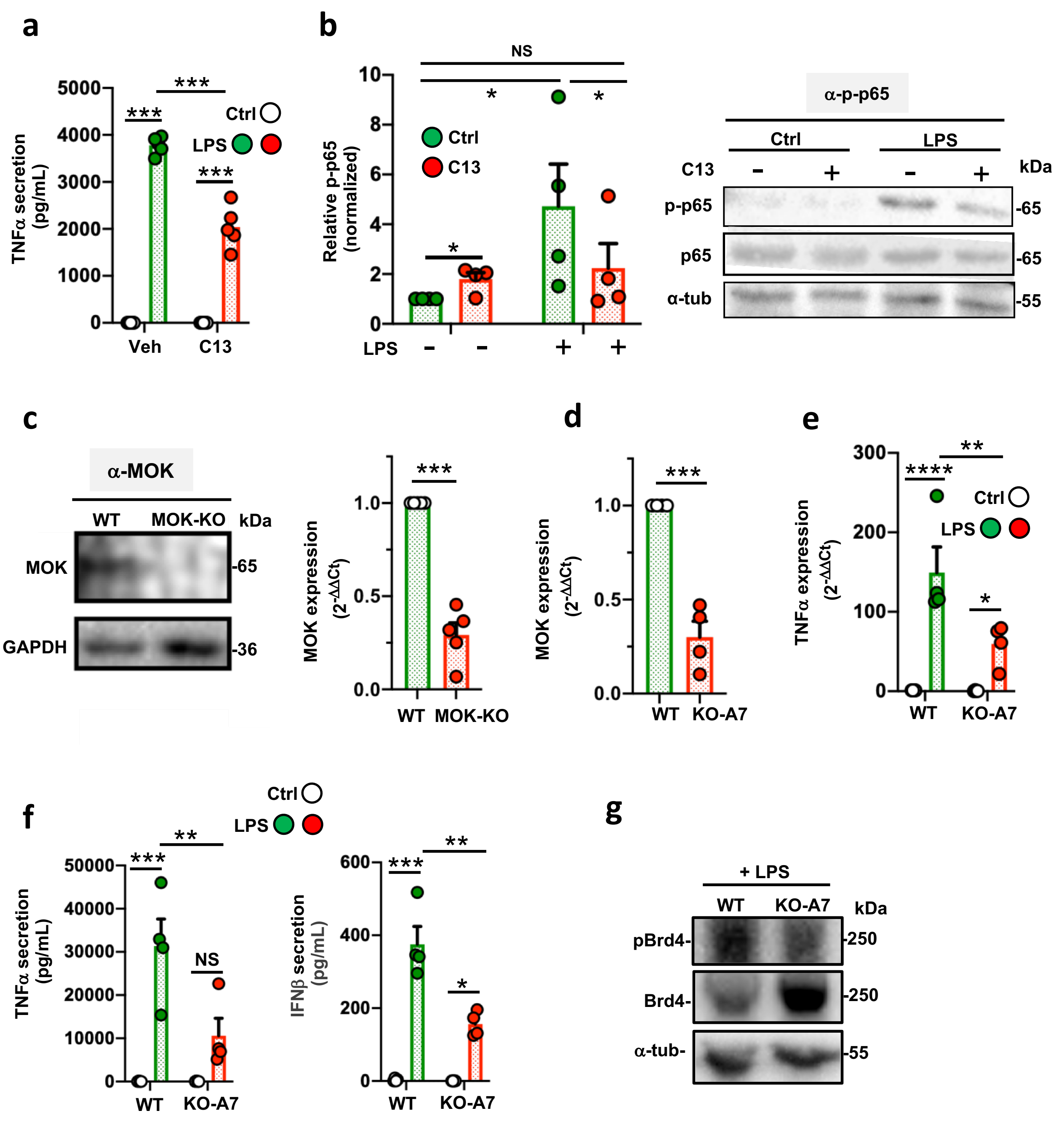


**Supplem. Figure 2: a)** Determination of TNFα secretion after stimulation of SIM-A9 cells with 1 μg/mL LPS for 5 h, pre-treated with 10 μM C13 (or vehicle) for 1 h. **b)** Quantification by band densitometry and representative image of Western blot assays using cell lysates from SIM-A9 cells stimulated or not with 1 μg/mL LPS for 5 h, and pre-treatment with 10 μM C13 (or DMSO) for 1 h, with anti-phospho-p65 (p-p65) and anti-p65 antibodies. Data represent means -/+S.E.M. normalized to untreated control (N=4). Student's t-test between shown groups, ratio-paired, two-tailed. **c)** Assessment of MOK expression in WT SIM-A9 cells and CRISPR/Cas9-generated MOK-KO clones by Western blot (left) and qRT-PCR (right). A band of ca. 65 kDa MW, in agreement with previoulsy reported for brain tissue (34), was observed in WT and not in MOK-KO cells. **d)** Assessment of MOK expression in WT SIM-A9 cells and MOK-KO A7 clone (KO-A7) by qRT-PCR. **e,f)** Determination of cytokine expression after stimulation with 1 μg/mL LPS for 5 h by qRT-PCR **(e)** or by ELISA **(f)** in KO-A7 cell clone. **g)** Representative image of Western blot assay from two independent experiments, using anti-pBrd4, anti-Brd4 or anti-α-tubulin, with lysates of WT and KO-A7 cells stimulated with 1 μg/mL LPS for 1 h.

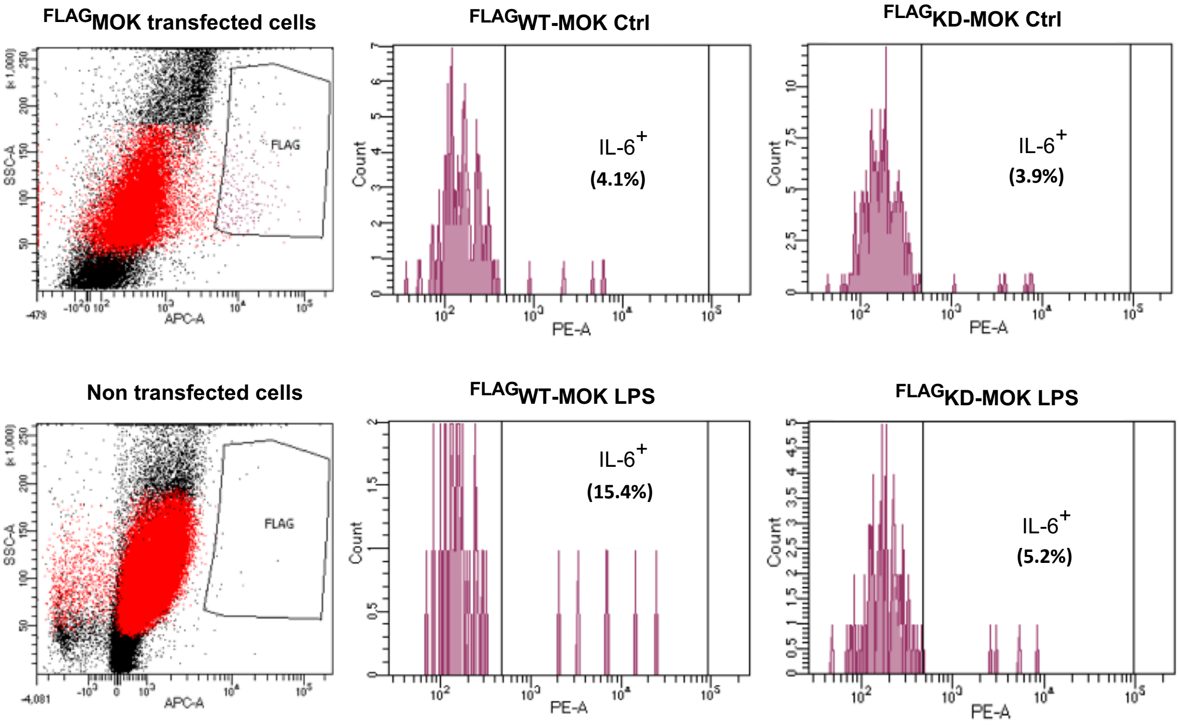


**Supplem. Figure 3:** Flow cytometry analyses for IL-6 expression (with PE-anti-IL-6 antibody) of MOK-KO cells transfected with $_{FLAG}$WT-MOK or $_{FLAG}$KD-MOK, stimulated or not with 1 μg/mL LPS. Histograms show the FLAG$^{+}$ cell population (detected with APC-anti-FLAG antibody), gated for the region shown in the dot plots.

**a**

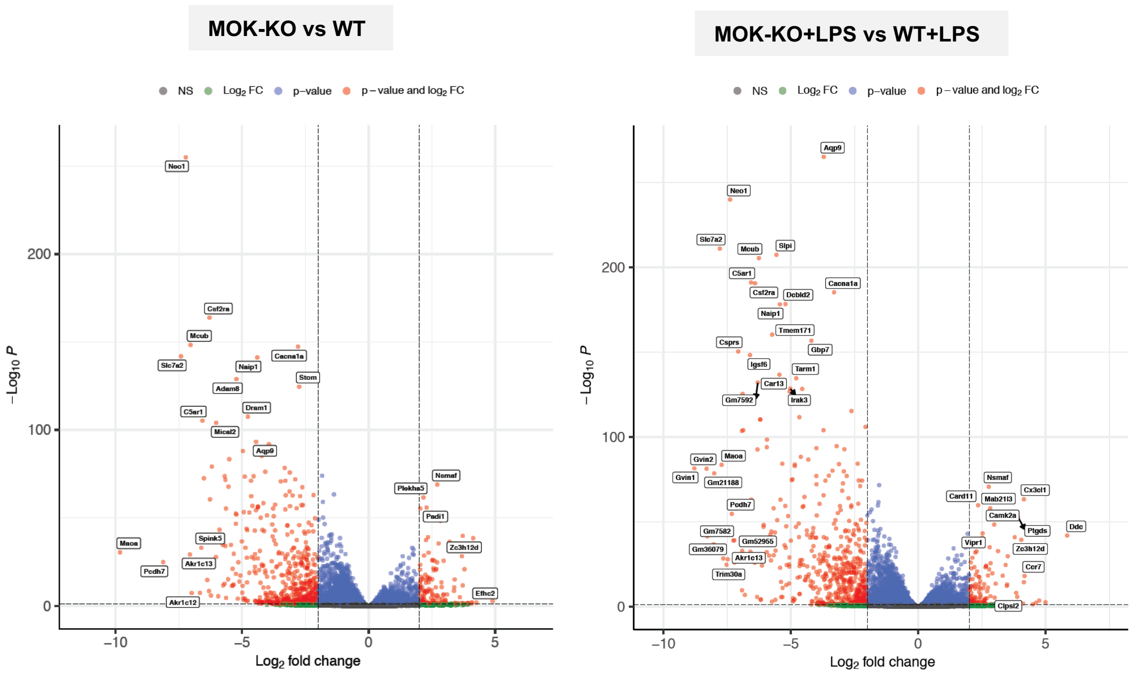


**b**

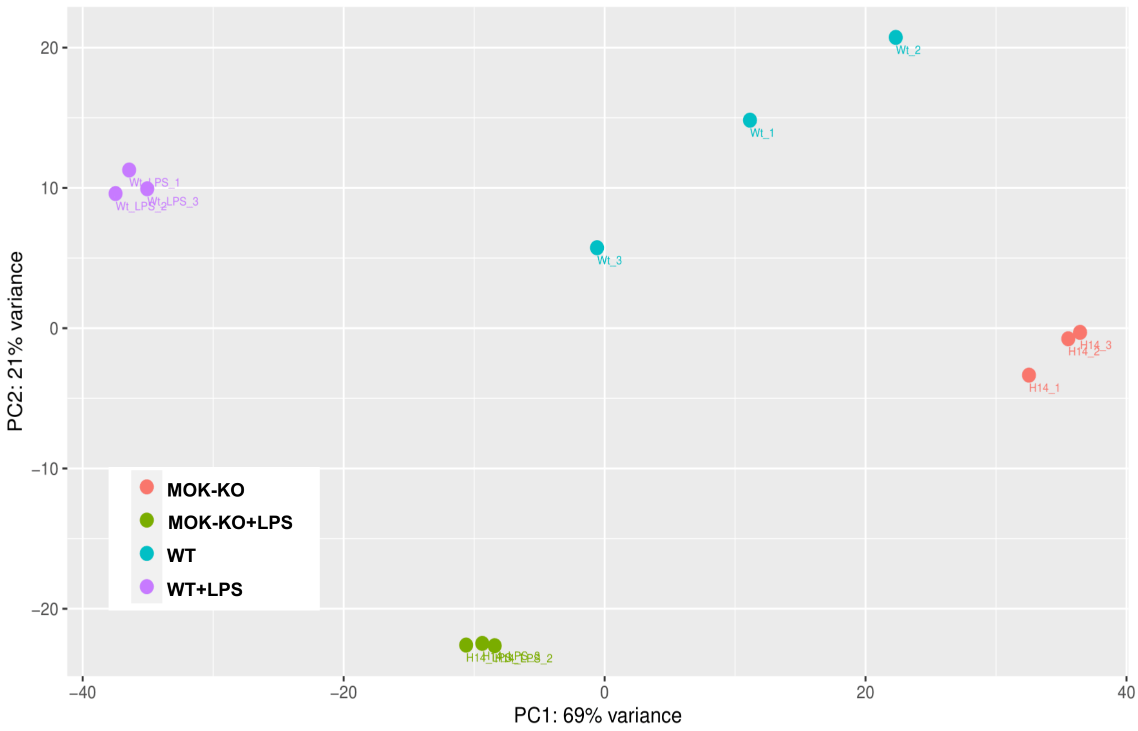


**Supplem. Figure 4:** RNA-Seq analysis from MOK-KO and WT SIM-A9 cells incubated (or not) with 1 µg/mL LPS for 5 h. **a)** Volcano plots representing all 45661 identified genes for MOK-KO vs. WT in the absence (left) and presence (right) of LPS-stimulation. pAdj<0.05 was considered significant. A few DEGs with the lowest p-values and largest fold-changes are labeled. **b)** PCA plot of RNA-Seq results (shown in Figure 3). Data are from three independent experiments (N=3).

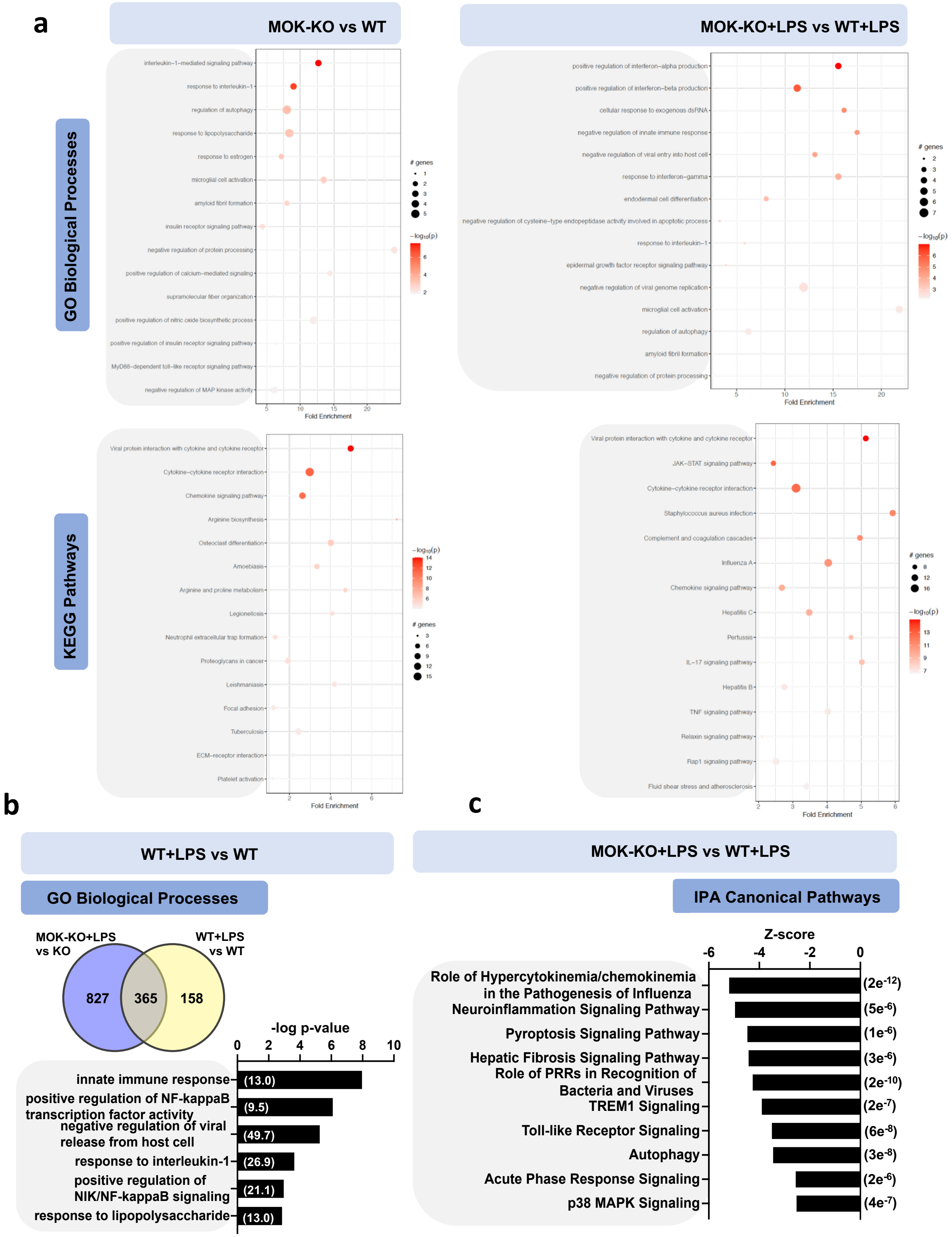


**Supplem. Figure 5:** RNA-Seq analysis from MOK-KO and WT SIM-A9 cells incubated (or not) with 1 μg/mL LPS for 5 h. Results are from three independent experiments (N=3). **a)** Top-15 enriched terms of GO 'biological processes' and KEGG pathway enrichment analyses performed using PathfindR (v1.6.2) by using all DEGs after applying the filter on Log2 fold-change values at ±2 along with pAdj<0.05 as input. **b)** Top GO 'biological process' enriched terms between WT+LPS vs. WT from the 985 protein-coding DEGs which are not common to both groups (Venn diagrams, 827+158). Fold-enrichments are shown in parentheses. Shown are the top hits based on p-value with at least three up/downregulated genes. **c)** Top 10 IPA-identified pathways based on p-value (shown in parentheses) that were differentially regulated (in any comparison with pAdj<e-5) in MOK-KO +LPS compared to WT +LPS.

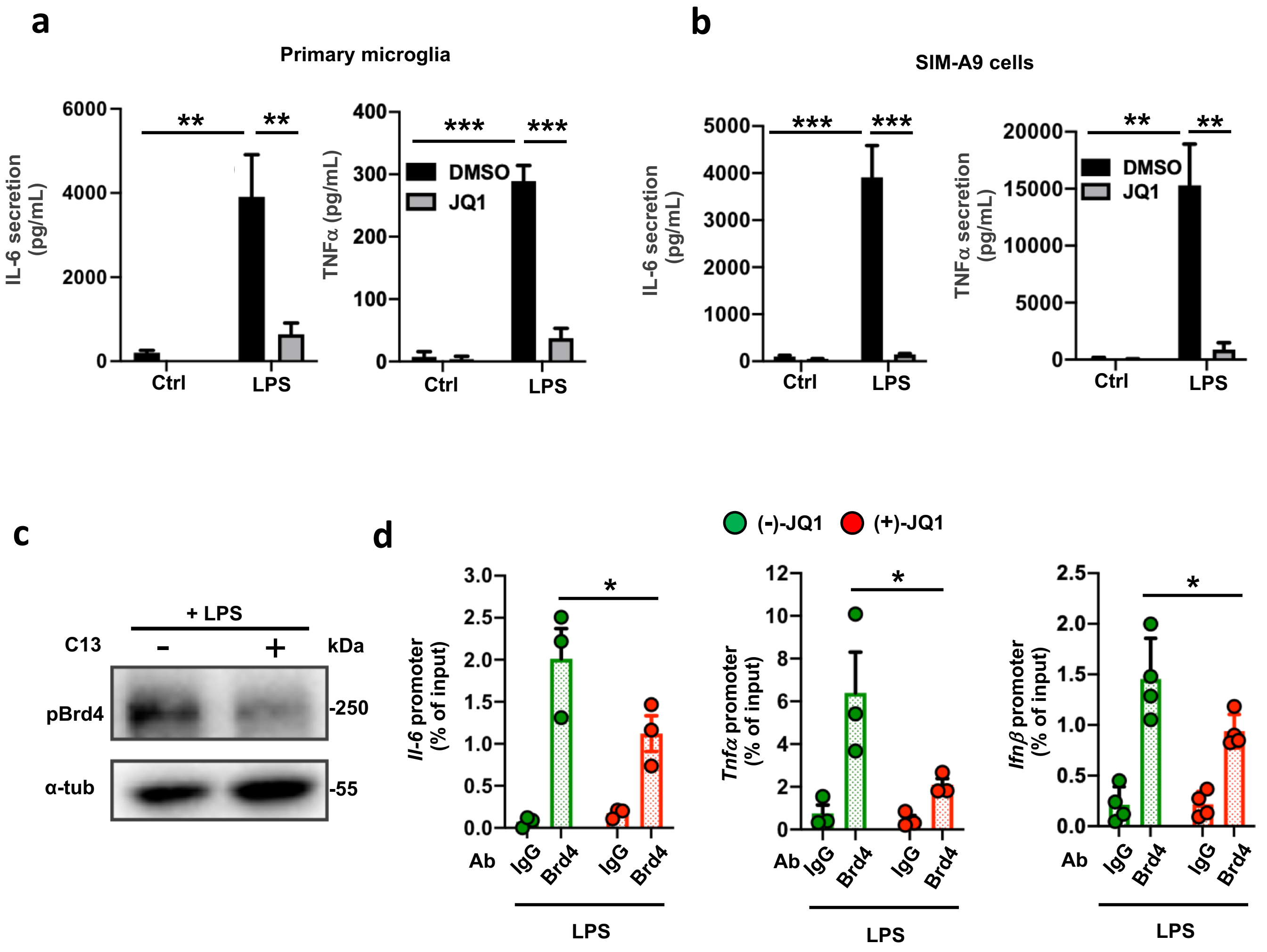


**Supplem. Figure 6.** Quantification of IL-6 and TNFα secretion by ELISA in supernatants from primary microglial **(a)** and SIM-A9 **(b)** cells pre-treated with 10 μM (+)-JQ1 (or DMSO) for 1 h and stimulated with 1 μg/mL LPS for 5 h (N=3). *P<0.05, **P<0.01, ***P<0.001. One-way ANOVA followed by Tukey post-hoc test. **c)** Western blot with cell lysates from 1 μg/mL LPS-stimulated WT SIM-A9 cells pre-treated or not with C13 (10 μg/mL, 1 h), using anti-pBrd4 or anti-α-tubulin antibody, as indicated. A representative image of two independent experiments is shown. **d)** ChIP-PCR assays from LPS-stimulated (1 μg/mL LPS, 1 h) WT SIM-A9 cells pre-treated with 10 μM (+)JQ1 or (-)JQ1 for 1h. Represented data are relative values (percentage of input) of PCR products for *Il6*, *Ifnb1* and *Tnfa* promoters after ChIP assay with either anti-Brd4 or IgG (control) antibodies. Student's t-test, unpaired, one-tailed. Data represent the mean percentages of input from 3 independent experiments (N=3) and error bars denote S.E.M.

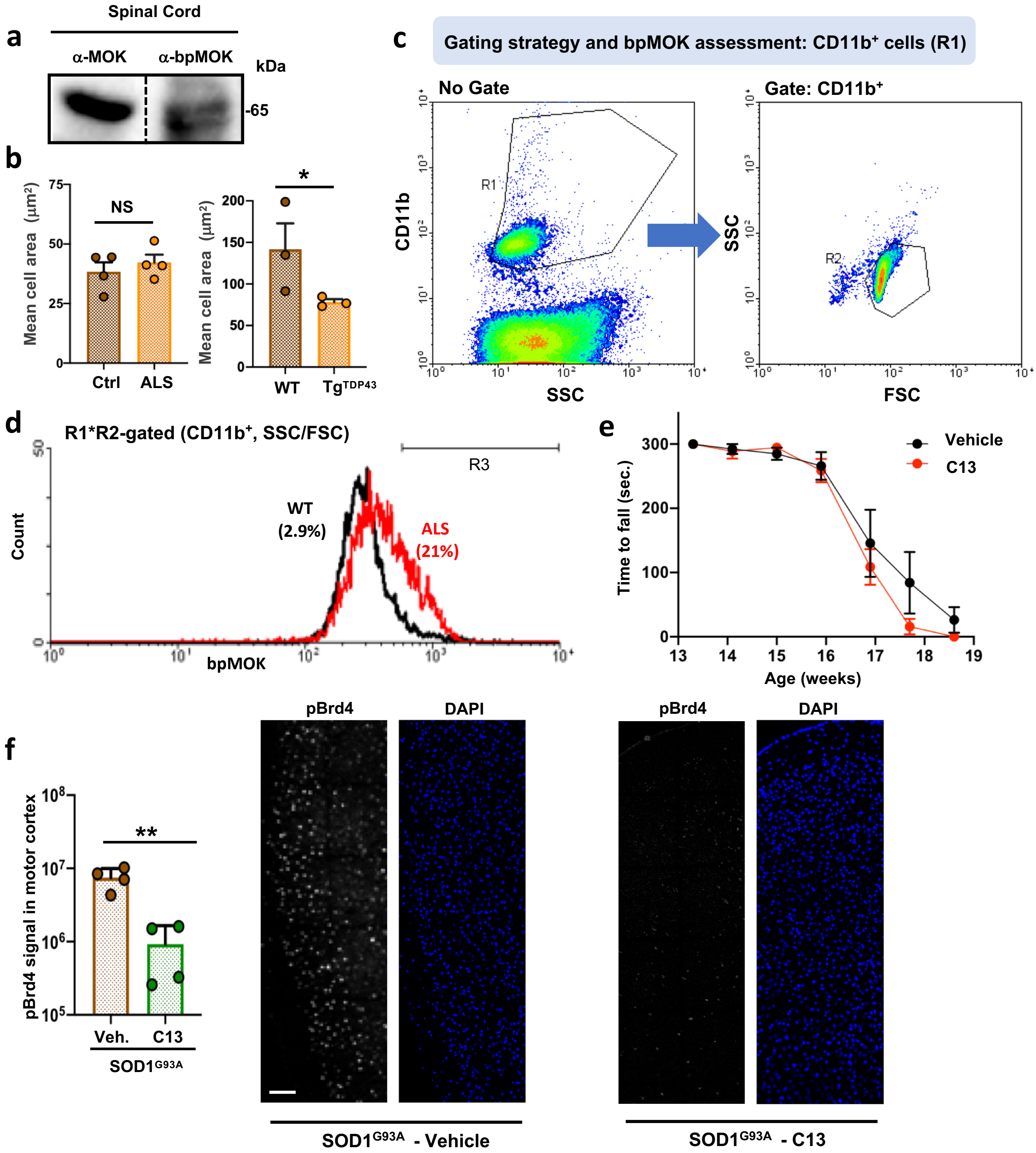


**Supplem. Figure 7. a)** WB of spinal cord tissue homogenates, blotted against anti-MOK or anti-bpMOK as indicated. **b)** Determination of cell size distribution by IHC analysis (with anti-bpMOK) of spinal cord tissue samples from Tg$^{TDP43}$ vs. WT mice (late stage; N=3) and from sporadic ALS patients vs. control subjects (N=4). Student's t-test, unpaired, one-tailed. **c,d)** Ficoll gradient-purified cells from 14-weeks old WT and SOD1$^{G93A}$ mouse spinal cords were immunolabeled with anti-CD11b and anti-bpMOK, and analyzed by flow cytometry with the gating strategy shown in **(c)**. The histograms shown for bpMOK fluorescence analysis of R2-gated (CD11b$^{+}$) cells correspond to one representative WT mouse (black) and one representative SOD1$^{G93A}$ mouse **(d)**. **e)** Motor performance assessment by rotarod test was carried out regularly since week 13 up to each mouse's end-point. Mean values and SEM are represented (N=6). Student's t-test, unpaired, one-tailed. **f)** Quantification of pBrd4 levels (integrated density) by IF with anti-pBrd4 of brain motor cortex samples from SOD1$^{G93A}$ mice after C13- or vehicle-treatment (week 15; 4 mice per experimental group, N=4, left). Representative IF images with DAPI counterstaining are shown (right). Scale bar: 100 μm. Student's t-test, unpaired, two-tailed. Data represent mean -/+S.E.M.

**Supplemental Materials and Methods**

**Animal models.** For IHC studies, spinal cord tissue slices (lumbar, sacral) were from late-stage Prp-TDP43$^{A315T}$ mice (Tg$^{TDP43}$), an ALS/FTD model described previously (35), and wild-type, age-matched controls. For all the remaining studies, transgenic B6SJL-TgN(SOD1-G93A) 1Gur/J (Jackson Laboratory, USA) male mice carrying the human SOD1 gene with the G93A point mutation (SOD1$^{G93A}$) (77) and wild-type counterpart mice were used. They were maintained at CABIMER animal facility on a 12-hour light/dark cycle, housed at 4-8 animals/cage with constant temperature (22 °C) and humidity (60%), and with food and water available *ad libitum*. Genotyping to assess mutant SOD1 copy number was performed for every mouse used in experiments. At time of weaning, littermates were identified through ear punching and separated in different cages according to their gender. The ear tissue extracted during the tagging procedure was then used to run a qRT-PCR for genotyping of the progeny. Animals were anesthetized intraperitoneally by injection of ketamine (100 mg/kg body weight) and xylazine (10 mg/kg body weight). Experiments followed the approved institutional animal protocols at the CABIMER institute for the care and use of laboratory animals, with ethical committee numbers 25-2010 and 02-2016 and complied with national and European Union legislation (Spanish RD 53/2013 and EU Directive 2010/63) for the protection of animals used for scientific purposes.

**Acute microglial cell isolation from adult mice.** Microglial cells from adult mice were acutely isolated from spinal cord tissues of 14 weeks-old mSOD1$^{G93A}$ Tg mice and their WT littermates (N=8), or from C13- or vehicle-treated mSOD1$^{G93A}$ Tg mice (N=4) and their WT littermates (N=6), accordingly. Mice were anesthetized by intraperitoneal injection of a Ketamine/Xylazine mixture, and intracardially perfused with phosphate-buffer saline (PBS). Spinal cords were extracted by flushing the spinal canal with PBS and minced into 2-3 mm pieces with a disposable scalpel in ice-cold Hank´s-balanced salt solutions (HBSS, cat. 14170-088). Each spinal cord microglial preparation was derived from a single animal by enzymatic digestion and mechanical disruption, and separation by density gradient, as detailed. Spinal cord fragments were dissociated with 0.25 % trypsin-EDTA for 10 min at 37 °C (GIBCO, cat. 25200072). Then trypsin treatment was stopped by adding RPMI 1640 medium (Sigma, cat. R8758) with 10% (v/v) FBS. The spinal fragments were then mechanically dissociated by repeated pipetting and by passing through a 70 μm cell strainer (Falcon cat. 352350s). Microglia was further purified for myelin removal by using Percoll gradient. Cells were collected by centrifugation at 350g for 10 min at 4 °C, the cell pellet resuspended in 2 ml of 70% isotonic Percoll, and then carefully over layered with 3 mL of 35% Percoll in PBS, and 2 ml of PBS. The discontinuous gradients were centrifuged at 850 g for 35 min at room temperature. The upper layer containing the white in appearance myelin debris was then removed. Cells were isolated from the gradient interface, washed in three volumes of PBS by centrifugation at 500 g for 5 min, and resuspended in RPMI 1640 medium supplemented with 2% FBS for cell counting and flow cytometry analysis.

**Immunolabeling of acutely isolated microglial cells and flow cytometry analysis.** Cells were stained on ice for 30 min with rat anti-Cd11b-APC (BD Biosciences, cat. 553312) or appropriate antibody isotype controls Rat Anti-Mouse IgG1-PE (BD Biosciences, cat. 562027), diluted in PBS with 1% FBS and 1mM sodium azide. For intracellular labelling of bi-phosphorylated MOK protein (bpMOK), cells were fixed and permeabilized with the Cytofix/Cytoperm Plus Fixation/Permeabilization Kit (BD Biosciences, cat. 554722) according to the manufacturer instructions and stained with anti-bpMOK antibody (pThr159+pTyr161, StressMarq, cat. SPC-1030; see Table S1 for antibody information), followed by donkey anti-rabbit IgG-Alexa Fluor 488 (Invitrogen, cat. A-21206). Antibodies for intracellular labelling were diluted in Perm/wash Buffer (BD Biosciences, cat. 554723). Samples were analysed using a FACSCalibur (BD Biosciences, Madrid, Spain). Data were analysed using either Cell Quest Software (Becton Dickinson, Mountain View, CA, USA) or the WinMDI 2.8 software (from http://www.cyto.purdue.edu/flowcyt/software/Winmdi.htm).

**Immunohistochemistry (IHC) and IHC analyses.** IHC assays and quantification were double blinded. Fixed spinal cord segments from $Tg^{TDP-43}$ and control mice and controls (lumbar region) or human sporadic ALS patients (with no known mutations, lower thoracic or higher lumbar region and obtained as explained before), were embedded in paraffin and sectioned to 10 μm-thick slices. After deparaffinization, sections were incubated with 1x citrate buffer (Cat. C9999, Sigma Aldrich) for 30 min at 95 ºC for antigen retrieval, followed by incubation in 3% (w/v) $H_2O_2$ to inhibit endogenous peroxidase activity. Sections were then incubated with 1% Bovine Serum Albumin (BSA) blocking solution for 1 h at room temperature, followed by overnight incubation at 4 ºC with anti-bpMOK antibody (pThr159+pTyr161, StressMarq, cat. SPC-1030). Then, sections were washed and incubated with the appropriate secondary biotinylated antibodies, followed by incubation with the avidin-biotin complex (Vectastain ABC-HRP Kit, Peroxidase-Rabbit IgG; Vector Laboratories, USA). Finally, sections were treated with diaminobenzidine-nickel substrate (DAB-nickel; Vector Laboratories) and then examined under a Leica DM6000B microscope. Images were acquired at 40x magnification. For comparative signal quantification, cell size and signal intensity for at least 50 cells (with visible nucleus on the plane) from three or four different, non-overlapping fields were assessed for each sample (corresponding to each individual) with the Fiji software (OpenSource). The background intensity -determined from 5 regions from 4 different images- was subtracted from the calculated signal intensities. In human samples, the less abundant very large cells (>2000 pixels, corresponding to >110 $\mu^2$) were excluded from the analysis. The mean intensity per cell and the intensity per cell area were calculated from all analysed cells for each sample/individual.

**Immunofluorescence (IF) of cultured cells.** For IF analyses, SIM-A9 cells or highly pure primary microglial cultures obtained as described above were used. Primary cell cultures were seeded on pre-treated poly-D-Lys (Sigma-Aldrich) glass lamellas. For the immunolabeling step, treated microglial cultures were fixed in cold PBS that contained 4% paraformaldehyde (Sigma-Aldrich) for 20 min at 4°C. After washing with PBS, cells were incubated with PBS that contained 0.5% (v/v) Triton X-100 (Sigma-Aldrich) and 3% bovine serum albumin (Sigma-Aldrich) for 1 h at 4 °C, for blocking and permeabilization. Cells were incubated overnight at 4 °C with Phospho-Brd4 (Millipore, Cat. ABE1451) primary antibodies (1:500). After washing with PBS, cells were incubated for 1 h at room temperature with their corresponding conjugated secondary antibodies, which was Alexa-Fluor488 donkey anti-rabbit IgG (Thermo Fisher, Cat. A-21206) (1:1000). After washes with PBS, nuclei were counterstained by incubating cells with PBS that contained 1 μg/mL DAPI (Sigma-Aldrich, Cat. D9542) for 15 min at room temperature. Fluorescence images were captured with confocal automated inverted microscope (Confocal Lase Scanning Microscope TCS SP5; Leica Microsystems). Images were taken at 63x magnification from randomly chosen fields.

**Immunofluorescence (IF) of spinal organotypic culture and brain slices.** Following treatment with LPS (or medium alone), spinal cord organotypic slices were fixed with

4% paraformaldehyde in PBS for 3 h at 4 °C. After blocking with 5% normal donkey serum (Merck Millipore) and 0.5% (v/v) Triton X-100 in PBS, sections were incubated with anti-bpMOK or anti-pBrd4 antibodies from rabbit. Slices were thoroughly washed in PBS with 0.2% Tween-20 and incubated with Alexa-Fluor 488-labeled donkey anti-rabbit IgG secondary antibody (1:1000; Thermo Fisher Scientific) diluted in PBS with 0.1% Triton X-100 and 1% of normal donkey serum for 2 h at room temperature. Finally, cell nuclei were labelled with 1 μg/ml DAPI for 15 min in PBS and sections were mounted with Vectashield medium (Vector Laboratories, Burlingame, CA, USA). Slices were analysed under confocal microscope (Confocal Laser Scanning Microscope TCS SP5; Leica Microsystems).

Removed murine brains were embedded in paraffin, stored at 4 ºC and then sectioned to 7 μm-thick slices. After deparaffinization, sections were incubated with 1x citrate buffer (Cat. C9999 Sigma Aldrich) for 30 min at 95 ºC for antigen retrieval, and washed three times with 0,1% (v/v) Triton in PBS (PBS-triton) during 5 minutes, followed by incubation in 3% (w/v) $H_2O_2$ to inhibit endogenous peroxidase activity. Sections were then incubated with 1% BSA and 3% normal donkey serum diluted in PBS-triton (blocking solution) for 1 h at room temperature and were stained overnight at 4 ºC with Phospho-Brd4 (Millipore, Cat. ABE1451) primary antibodies (1:100) diluted in blocking solution. After washing with PBS-triton, cells were incubated for 1 h at room temperature with their corresponding conjugated secondary antibodies diluted in blocking solution, which were Alexa-Fluor546 donkey anti-rabbit IgG (Thermo Fisher, Cat. A-10040) (1:500). After washes with PBS-triton, nuclei were counterstained by incubating cells with blocking solution that contained 1 μg/mL DAPI (Sigma-Aldrich, Cat. D9542) for 30 min at room temperature and sections were mounted with Vectashield medium (Vector Laboratories). Fluorescence images of complete brain slices were captured with automated inverted microscope (DMi8 S; Leica Microsystems) at 40x magnification. For quantification, the same region of interest (ROI) was manually selected belonging to the motor cortex of the brains. ImageJ software was used to measure the integrated density of the fluorescence of these ROIs.

**Cytokines determination by ELISA.** Primary microglia or SIM-A9 cell cultures were stimulated as described before. Harvested culture supernatants were centrifuged at 400 g for 5 min, and cell-cleared supernatants were recovered and stored at -20 °C before cytokine measurement. IL-6 and TNF-α levels were assayed with Mouse IL-6 and TNF-α BD OptEIA ELISA sets (BD Biosciences, Cat. 555240 and 555268, respectively). IFN-β and IL-1α were assayed with Mouse IFN-β and IL-1α/IL-1F1 DuoSet ELISA kits (R&D Systems, Cat. DY8234 and DY400, respectively). All cytokine quantifications were done by following the manufacturer's protocol.

**Transfection of microglial cell cultures**. For the overexpression of FLAG-tagged WT-MOK and KD-MOK constructs in SIM-A9 cells, Glial Mag transfection reagent (OzBiosciences, Cat. KGL0250) was used. The protocol was carried out according to manufacturer's instructions. Plasmids were diluted in Optimem medium (Gibco, Cat. 31985062), and 1 μL of Glial Mag and 0.5 μg of plasmid were employed and volumes and concentrations were optimized for a 24-well plate assay. Transfection time was of 30 h in total, including a 5 h-stimulation with or without 1μg/mL LPS or GolgiStop (BD Biosciences, Cat. 554724).

**Immunolabelling and flow cytometry of transfected cells**. SIM-A9 cells were fixed and permeabilized for 20 min at 4 ºC by using Fixation/Permeabilization solution (BD Biosciences, Cat. 555028). After washing twice with Wash Buffer (2 mM EDTA, 2% v/v FBS in PBS), cells were incubated for 15 min at room temperature in Perm/Wash Buffer (BD Biosciences, Cat. 555028). Immunostaining was performed using APC-conjugated anti-FLAG (Abcam, Cat. Ab72569) and PE-conjugated anti-IL-6 (BD Biosciences, Cat. AB395367) at a 1:40 dilution in both cases in Perm/Wash Buffer, and samples were incubated at 4 ºC in the dark for 30 min. After antibody incubation, cells were washed twice with Perm/Wash Buffer and resuspended in Wash Buffer, and were kept at 4 ºC in the dark until analyses with LSR Fortessa X-20 cytometer (BD Biosciences).

**Preparation of cell lysates**. Cells were cultured in 12-well plates at 400,000 cells/mL for 24 h previous to treatment. After treating as required, cells were washed with cold PBS and lysed in lysis buffer (50 mM Tris pH 7.4, 150 mM NaCl, 0.25% deoxycholate, 10% glycerol, 0.1% SDS, 0.01% IGEPAL) containing Complete EDTA-free Protease inhibitor cocktail tablets (Roche, Cat. 11836170001), 1 mM PMSF (ITW Reagents, Cat. A0999) and PhosSTOP (Roche, Cat. 04906845001). Lysates were incubated for 5 min at 37 ºC followed by a 20 min incubation in ice with manual agitation every 5 min They were then centrifuged at 12,000 g for 5 min at 4 °C. Cleared supernatants were collected and stored at -20 °C until use. Protein concentration was determined using Micro BCA™ Protein Assay Kit (Thermo Fisher, Cat. 23235), following manufacturer's instructions.

**Western blot assays**. Equal volumes of cell lysate (20 μl) with comparable protein content (40-50 μg) were subjected to 10% or 12% SDS-PAGE electrophoresis and semi-dry or wet transfer to a 0.2 μm nitrocellulose or PVDF membrane (Amersham, Cat. 10600006 and Amersham, Cat. 10600021, respectively). After blocking with 5% (w/v) non-fat milk in Tris-buffered saline with 0.05% Tween 20 (TBST) during 1h at room temperature (RT) and washings with TBST, membranes were incubated overnight at 4 ºC with the corresponding primary antibodies against: (Ser492) Phospho-Brd4 (Millipore, Cat. ABE1451) (1:500), Brd4 (Bethyl, Cat. A301-985A100) (1:1000), MOK (Abbexa, Cat. abx129598) (1:1000), Ser536-Phospho-NFκB p65 (ThermoScientific, Cat. PA5-17782) (1:1000), p65 (Cell Signalling, Cat. 3034) (1:1000). The following day membranes were washed with TBST and incubated with the appropriate HRP-conjugated secondary antibody: goat anti-mouse IgG (Pierce, Cat. 32430) (1:2500) or goat anti-rabbit IgG (Pierce, Cat 31460) (1:10000), for 1 h at room temperature. Finally, after washings with TBST, the substrate solution ECL reagent (Merck Millipore, Cat. WBKLS0500) was added and bands were detected using a ChemiDoc™ MP Imaging System (Bio-Rad, Cat. 1708280). The results were quantified using the ImageLab Software 6.1 and normalized against the housekeeping gene GAPDH (Millipore, Cat. MAB-374) (1:2500) or α-tubulin (Sigma-Aldrich, Cat. T6199) (1:6500), accordingly.

**Cell lysate preparation for anti-phospho-Ser/Thr immunoprecipitation.** After treatment of primary microglial cells with TDP-43 or sham aggregates as described before, culture supernatants were harvested. Following two washes with PBS, cells were lysed with NP-40 lysis buffer (Invitrogen, FNN0021) supplemented with protease inhibitors cocktail (Roche), 1 mM PMSF (ITW Reagents, Cat. A0999) and PhosSTOP (Roche, Cat. 04906845001). For the IP procedure, Protein A Dynabeads (Invitrogen, Cat. 10001D) were used. After incubating the beads with 2 μg of anti-phospho-Ser/Thr rabbit antibody (Abcam, Cat. ab17464) for 1h at room temperature under rotation, crosslinking of the captured antibody was performed with BS3 (Thermo Fisher, Cat. 21580) according to the manufacturer's protocol. Afterwards, the beads-antibody conjugates were added to the cell lysates and incubated for 1 h at room temperature under rotation. After washing, elution was carried out with 50 mM glycine solution (pH 2.8) and the eluate was pH-equilibrated with PBS for subsequent LC-MS/MS analysis.

**LC-MS/MS and protein identification.** IP protein eluates were subjected to TCA/acetone precipitation and the pellet was resuspended in a solution of 0.2% RapiGest in 50 mM ammonium bicarbonate. DTT was added to 5 mM final concentration and incubated for 30 min at 60 ºC. Afterwards, chloroacetamide was added to 10 mM final concentration and incubated for 30 min at room temperature in the dark. Trypsin digestion was done overnight at 37 ºC at a 1:40 (trypsin:protein) ratio. The next day, trypsin was inactivated with formic acid and acetonitrile was added to a final concentration of 2%. Injection volume was 10 µL per run.

LC-MS/MS analyses were done in a TOF (5600 Plus, Sciex) triple quadrupole equipped with a nano electrospray source coupled to a nanoHPLC Eksigent. Analyst TF 1.7 software was used for control of equipment and for the acquisition and processing of data. Peptides were first loaded in a 'trap column' (Acclaim PepMap 100 C18, 5 µm, 100 Å, 100 µm id × 20 mm, Thermo Fisher Scientific) in an isocratic manner in 0.1% formic acid/2% acetonitrile (V/V) at a 3 µL/min flow for 10 min Then, they were eluted in a reverse-phase analytical column (Acclaim PepMap 100 C18, 3 µm, 100 Å, 75 µm id × 150 mm, Thermo Fisher Scientific) coupled to a 'PicoTip emitter' (F360-20-10-N-20_C12, New Objective). Peptides were eluted with a linear gradient of 2-35 % (v/v) solvent B in 60 min at a 300 nL/min flow. Solvents A and B were 0.1% formic acid (v/v) and acetonitrile with 0.1% formic acid (v/v), respectively. Source voltage was selected at 2600 V and heater temperature was maintained at 100 ºC. Gas 1 was selected at 15 psi, gas 2 at zero, and curtain gas at 25 psi. For protein identification, acquisition was carried out with a Data Dependent Acquisition (DDA) method consisting of a TOF-MS with a sweep window of 400-1250 m/z, accumulation time of 250 ms, flowed by 50 MS/MS with a sweep window of 230-1500 m/z, accumulation time of 65 ms and a total cycle time of 3.54 s.

For protein identification, the ProteinPilot v5.0.1 (Sciex) software was used with a Paragon method, with trypsin as the enzyme and iodoacetamide (IAA) as the Cys alchylating agent, using the reference proteome of mouse (*mus musculus*) downloaded from uniprot.org (02/2021) in a FASTA format as the databank, fused with the Sciex contaminants. A false-positives analysis was done and proteins with FDR (False Discovery Rate) lower than 1% were selected.

**RNA isolation and cDNA synthesis for qRT-PCR.** Total RNA from SIM-A9 or primary microglial cells was extracted by using Primezol (Canvax, Cat. AN1100). Cells were directly lysed in 12-well culture plates by adding 1 mL of Primezol per well. Samples were incubated 5 min at room temperature and then 200 µL of chloroform was added. Tubes were shaken vigorously by hand for 15 sec. and incubated 3 min at room temperature and then centrifuged at 12000 g for 15 min at 4 ºC. The upper layer containing RNA was separated and transferred into a new tube without disturbing the interphase. RNA was precipitated by adding 0.5 mL of isopropyl alcohol and incubated for 10 min at room temperature. Tubes were centrifuged again at 12000 g for 10 min at 4 ºC. Supernatant was removed and pellet was washed once with 70% ethanol. Finally, tubes were centrifuged at 7500 g for 5 min at 4 ºC. Supernatant was removed once more and pellet was air-dried. Pellet was dissolved in RNAse-free water and stored at -80 ºC. After extraction, 1 µg of RNA was reverse-transcribed by using PrimeScript RT reagent Kit (Takara, Cat. RR047Q) following the manufacturer's guidelines.

**qRT-PCR.** Quantitative real-time PCR (qRT-PCR) with template cDNA was performed by using iTaq Universal SYBR Green Supermix (Bio-Rad, Cat. 1725124) on a QuantStudio 5 Real-Time PCR System. Primers pairs (Sigma-Aldrich) were preferentially designed or selected to anneal in different exons and they were as follows: MOK, HPRT, IL-6, IFNβ, TNFα, IL-1α, IRF7 (Table S3). Amplicons were analysed for 40 cycles by using an optimized qRT-PCR thermal profile. Changes in gene expression were determined by applying the formula $2^{-\Delta\Delta Ct}$ where ΔCt is calculated taking HPRT as endogenous control. ΔΔCt values were calculated by subtracting the average ΔCt values obtained for non-treated cell samples.

**RNA-Seq analyses.** RNA-Seq analyses were carried out by the Genomics Unit, CABIMER (Andalusian Center for Molecular Biology and Regenerative Medicine, Seville, Spain). For transcriptional analyses of MOK-KO vs. WT SIM-A9 cells, total RNA-Seq was used. After removing the culture supernatant, cells were washed twice with PBS and RNA was extracted using the RNeasy Mini Kit (Qiagen, Cat. 74104) according to the manufacturer's guidelines. RNA samples were quantified using Qubit RNA HS Assay Kit and RNA size profile and integrity were analysed on Bioanalyzer 2100 (Agilent). RNA quality was considered 'excellent' and RNA integrity number estimates for all samples were between 9.7 and 10.0. After ribosomal RNA depletion, strand-specific (cDNA) libraries preparation was performed by applying Stranded TOTAL RNA prep Ligation with RIBO-ZERO PLUS kit (Illumina), following the manufacturer's instructions. Adapters (Illumina) included a sample/library-specific index. After library amplification by PCR and all recommended quality control assays, libraries were normalised using Qubit and quality control was performed with Bioanalyzer 2100. Individual libraries were pooled together at equimolar ratios and diluted to ~1.2 nM. Pooled libraries were denatured and further diluted prior to loading on the sequencer. Paired-end sequencing was performed using a NovaSeq6000 SP and 2x50bp (Illumina), generating a raw read count of >68 million reads per sample. Sequencing quality as assessed with BaseSpace Hub software (Illumina) was high (89.8% >Q30 and >106 Gb depth).

For mRNA-Seq transcriptional analyses from primary microglial cells, sequencing of mRNA was carried out. The procedure was as described above, with the following changes: mRNA was isolated by using poly-dT-coated beads and libraries were prepared by the TruSeq Stranded mRNA kit (Illumina). After library amplification by PCR and all recommended quality control assays, individual libraries were pooled together at equimolar ratios and diluted to ~1.4 pM. Paired-end sequencing was performed using NextSeq 500 Mid Output and 2x75pb (Illumina), generating a raw read count of >24 million reads per sample. Sequencing quality as assessed with BaseSpace Onsite software (v. 3.22.91.158, Illumina) was high (88% >Q30 and >24 Gb depth).

**RNA-Seq and mRNA-Seq data processing.** RNA-Seq sequencing data were generated from four cell treatments with three biological replicates each (12 samples in total), by using the paired end approach of Illumina technique. Total paired end fastq files were used for the RNA-Seq analysis via the pipeline: FastQC-Trimmomatic-STAR-FeatureCounts-DESeq2. STAR aligner (v. 2.7.9a) was used to perform read alignment and mapping to the reference genome, the latest release of the mouse genome from Ensembl (*Mus musculus*) genome database. The feature Counts module of R-package Rsubread (v. 2.0.1) was used for extracting raw read counts for all the samples. These raw read counts were then used to perform the differential expression analysis by using the DESeq2 package (v. 1.32.0) in R.

mRNA-Seq sequencing data were generated from two cell treatments with three biological replicates each (6 samples in total) as described above, with the following changes: Fastq files were generated with the pipeline FASTQ Toolkit (v1.0.0, Illumina) with filtering, timming and de-multiplexing data. The reference genome was *Mus musculus* (UCSC mm10). Cufflinks Assembly & DE, BaseSpace Workflow (v. 2.1.0, Illumina) was used to perform the differential expression analysis.

**S4U-labeling and SLAM-Seq analysis.** After pre-treatment with 10 μg/mL polymixin B, SIM-A9 cells in culture were incubated with 10 μg/mL C13 (or DMSO) for 1 h. Then, one tenth volume of medium without serum containing TDP-43 or sham aggregates (5 μg/mL final) was added to each well and incubated for 2.5 hours. This was followed by addition of 4-thiouridine (S4U, Cayman Chemical, Cat. 16373) to a final concentration of 100 μM (found by us to retain 80% of cells viability by MTT assay after 2 h) and incubation for a further 90 min in the dark. After washing with PBS, total RNA was extracted with easy-BLUE (iNtRON Biotechnology) and protected from light during the procedure to prevent S-S cross-binding. 5 μg of RNA was used for SLAM-seq modifications, as previously described (81). Briefly, iodoacetamide (Sigma I1149) was conjugated to 4SU in a 50 mM pH 8.0 phosphate buffer in DMSO/Water (1:1), the reaction was quenched with DTT and RNA was reprecipitated using ethanol and NaOAc. Libraries were prepared using Quantseq 3' mRNA-Seq Library Prep Kit (Lexogen). SLAM-Seq analysis was done similar to previously reported (82).

**Table S1. List of primary antibodies used**

| Antibodies | Host | Manufacturer | Catalog number | Application |
|---|---|---|---|---|
| MOK | Rabbit | Abbexa | abx027945 | IHC, WB |
| MOK (pT159+pY161) (bpMOK) | Rabbit | StressMarq Biosciences | SPC-1030 | IF Flow Cytom |
| Phospho-Ser/Thr | Rabbit | Abcam | ab17464 | IP |
| Brd4 | Rabbit | Bethyl | A301-985A100 | ChIP |
| Phospho Brd4 (Ser492) | Rabbit | Millipore | ABE1451 | IF, WB |
| Non-immune IgG | Rabbit | Sigma-Aldrich | I5006 | ChIP, Co-IP |
| FLAG, APC-conjugated | Mouse | Abcam | ab72569 | Flow Cytom |
| IL-6, PE-conjugated | Rat | BD Biosciences | AB_395367 | Flow Cytom |
| IL-18 | Rabbit | Mybiosource | MBS559603 | Western blot |
| p65 | Rabbit | Cell Signaling | Cat. 3034 | WB |
| p-p65 | Rabbit | Thermo Scientific | Cat. PA5-17782 | WB |
| CD11b, APC-conjugated | Rat | BD Biosciences | 553312 | Flow Cytom |
| Caspase-1 | Mouse | Adipogen | AG-20B-0042-C100 | WB |
| GAPDH | Mouse | Millipore | MAB-374 | WB |
| $\alpha$-tubulin | Mouse | Sigma-Aldrich | T6199 | WB |

**Table S2. List of secondary antibodies used**

| Antibodies | Host | Manufacturer | Catalog number | Application |
|---|---|---|---|---|
| Anti-rabbit IgG, HRP conjugated | Goat | Thermo Scientific (Pierce) | 31460 (1858415) | WB |
| Anti-mouse IgG, HRP conjugated | Goat | Thermo Scientific (Pierce) | 32430 (1858413) | WB |
| Anti-rabbit IgG, Alexa- Fluor488 | Donkey | Thermo Fisher | cat. A-21206 | IF |

**Table S3. List of PCR primers used**

| Primer name | Direction | Primer sequence |
|---|---|---|
| MOK | For | 5'-GTTCTACGAGATTGCCAGCCT -3' |
| | Rev | 5'-CTGCTTTCTGCCTTCCTGTGAGA -3' |
| HPRT | For | 5'-GGACTAATTATGGACAGGACTG -3' |
| | Rev | 5'-TCCAGCAGGTCAGCAAAGAA -3' |
| IL-6 | For | 5'-TCCGGAGAGGAGACTTCACA -3' |
| | Rev | 5'-TTCTGCAAGTGCATCATCGT -3' |
| TNFα | For | 5'-GCCTCTTCTCATTCCTGCTTG -3' |
| | Rev | 5'-CTGATGAGAGGGAGGCCATT -3' |
| IFNβ | For | 5'-CCACCAGCAGACAGTGTTTC -3' |
| | Rev | 5'-GAAGATCTCTGCTCGGACCA -3' |
| IL-1α | For | 5'-CGAAGACTACAGTTCTGCCATT -3' |
| | Rev | 5'-GACGTTTCAGAGGTTCTCAGAG -3' |
| IRF7 | For | 5'-GGTCGTAGGGATCTGGATGA -3' |
| | Rev | 5'-ACCTTATGCGGATCAACTGG -3' |
| IL6 prom. [28] | For | 5'-TGTGGGATTTTCCCATGAGT -3' |
| | Rev | 5'-TGCCTTCACTTACTTGCAGAGA -3' |
| TNFα prom. [71] | For | 5'-GGACTAGCCAGGAGGGAGAA -3' |
| | Rev | 5'-TGTCTTTTCTGGAGGGAGATGT -3' |
| IFNβ prom. [72] | For | 5'-GCCAGGAGCTTGAATAAAATG -3' |
| | Rev | 5'-CTGTCAAAGGCTGCAGTGAG -3' |
